# The representation of object concepts across the brain

**DOI:** 10.64898/2025.12.07.692815

**Authors:** Spencer R. Loggia, Helen E. Feibes, Stuart Duffield, Karthik Kasi, Bevil R. Conway

## Abstract

The mechanisms supporting concrete object concepts remain elusive. Here, macaque monkeys spent years learning object-concept primitives defined by color and shape. During fMRI, the monkeys held in mind one concept feature cued by the other. Using a whole-brain searchlight, we observed cross-feature decoding not in retinotopic cortex, but bilaterally in anterior inferior temporal cortex (aIT), temporal pole (TP), and frontal cortex (FC). Only TP and aIT had response patterns to concept shapes matching color-space geometry, and only PF had responses that decode behavioral errors. Two additional imaging experiments showed that color-associated versus non-color-associated shapes, and color-shape combinations congruent versus incongruent with the concepts, were distinguished only by extrastriate visual cortex, suggesting extrastriate cortex underpins color perception and perceptual feature binding. Together, the results provide a blueprint for how the brain generates stable object concepts and provides hypotheses for how conceptual representations engage perceptual representations through selective feedback during vision.

## Main

A stable concept is the “habitual or customary connection” between two ideas that is forged by the mind to enable adaptive behavior^1^. Concrete concepts correspond to tangible objects (e.g. banana, stop sign) and are thought to be used by the mind to construct abstract concepts (e.g. sustenance, traffic), providing building blocks for intelligence^2–4^. How concepts are formed and deployed therefore provides an important inroad to understanding the constructive nature of memory^5^. One widely recognized process by which concepts form is paired association, “the unconscious processes of the association of ideas,” as described by Hermann Helmholtz ^6^ and as instrumentalized by Mary Whitton Calkins^7^. Many object concepts link color and shape. The concept “banana” is prototypical: it associates one idea (yellow) with another idea (crescent) to predict a tasty treat. Such concepts are useful, as when searching for items in cluttered scenes where shape information required for object identification is occluded.

Where and how object concepts are represented in the brain is unresolved^8,9^. Among the possibilities are that they are grounded in sensory areas ^4,10–13^, mediated by an a-modal “hub” in anterior temporal lobes^14–16^, represented in prefrontal cortex^17^, or distributed broadly ^18–20^ (**Fig. 1a**). Arbitrating among the possibilities has been challenging for several reasons, including the difficulty in disentangling visual responses (which do not require memory) from conceptual responses (which do)^21^. For example, the neural representation of the concept of a familiar person cannot be recovered simply by comparing responses to photographs of that person and unfamiliar people because the images will differ in both conceptual content and visual features.

**Fig. 1:**
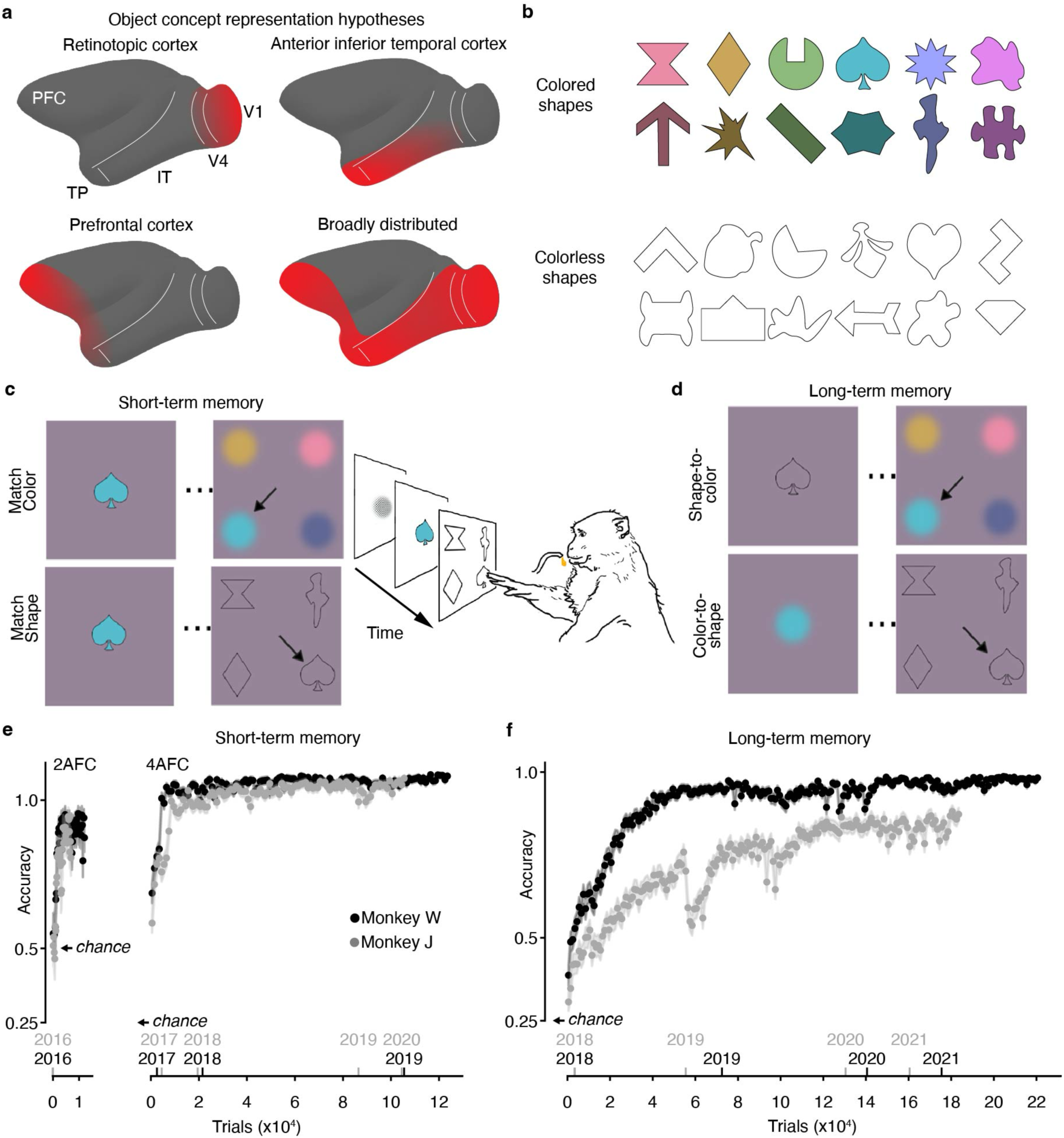
Learning object concepts defined by color and shape. a, Four hypotheses for how object concepts might be represented in the brain (see *Introduction*). **b,** The two sets of digital objects learned by the monkeys. **c,** The monkeys first learned about the colored objects in trials that could be completed with only short-term memory. The animals were shown a colored shape in one frame and in the next frame, they touched the matching shapeless color for “match color” trials (top) or the colorless shape for “match shape” trials (bottom). Trials were self-initiated and performed with a touchscreen tablet in the home environment. A juice drop was given for correct choices. The monkeys also performed trials in which the cue and match options were drawn from the set of colorless shapes (panel b, bottom). **d,** After the monkeys were successful in short-term memory trials, they were provided trials to reinforce long-term memory of the objects. They were shown a colorless color-associated shape and had to match it with a shapeless colored blob (“shape-to-color” trials, top), or a shapeless blob that they had to match with a colorless color-associated shape (“color-to-shape” trials, bottom). **e,** Learning curves on the short-term memory trials (solid symbols, each data point shows the accuracy of 200 trials for 2-alternative forced choice experiments (2AFC) and 1000 trials for 4AFC experiments; shading shows 95% CI generated by bootstrapping over trials per data point, chance is 0.5 (2AFC) or 0.25 (4AFC)). The x-axis shows both the cumulative number of trials and year timestamps. **f,** Learning curves on the long-term memory trials (conventions as **e**).

One way around this challenge is to test for responses to the associations that underwrite concepts. This approach has been used with microelectrode recordings to discover “pair-association” neurons in monkeys^22^ and “concept cells” in humans^23^. But microelectrode recording has limited sampling, making it difficult to determine how a concept repreparation is distributed across the brain. The possibility of broad distribution is perhaps expected since perception, stimulus familiarity, and memory are thought to engage many, if not all, the same brain regions^5,9,24–28^. Indeed, “concept cells” discovered in humans might not even be a necessary part of a concept representation since lesions of areas containing them do not ablate the concepts^29,30^.

Functional magnetic resonance imaging (fMRI) could provide the missing whole-brain perspective, but the technique has led to seemingly contradictory conclusions. Some studies suggest that perception and knowledge are grounded (or “embodied”) in a common neural substrate,^10,11,31–33^, with some studies suggesting that retinotopic cortex including V1 is reactivated when conceptual associations are called to mind^34–39^. But other studies implicate retinotopic visual cortex as an encoder of retinal data^40–42^ with no activation during mental imagery,^43,44^ leaving conceptual representations to higher-order areas^9,16^ such as prefrontal cortex^17,45^, a single spot in the left anterior temporal lobe^15,46^, or broadly in association cortex^18^.

The disparate conclusions may be explained by the fact that memory signals are not only small and hard to measure but also eclipsed by individual differences in brain organization and behavior. Indeed, a recognized roadblock to knowing where and how object concepts are represented is that the character, strength, and subjective value of concepts varies substantially among people^47,48^. Here, we overcome these obstacles with experiments in macaque monkeys, a model of human visual cognition. We provided the animals with lifetime experience of a set of simple color-shape concepts whose subjective value was fixed and whose physical features were precisely defined, providing a ground truth for the content of the conceptual representations.

Subsequent fMRI experiments in the animals show where the learned concepts are represented and provide clues to how the conceptual representations interact with perceptual representations through selective feedback during vision.

### Learning color-shape concepts

Two macaque monkeys acquired substantial experience with a set of 2-dimensional stimuli (**Fig. 1b**) through self-initiated interactions with a touch-screen tablet connected to a reward-delivery system in their home cage . We refer to the stimuli as “objects” and their neural representation as “object concepts.” Others have described these stimuli as “conceptual primitives”^49^ because they comprise the most basic concept unit, a paired association of behavioral relevance.

The controlled digital objects provide several advantages over natural objects for the present study. First, their shapes and colors were precisely controlled, so we are confident of the sensory ground truth that underwrites the resulting object concepts (in nature, color and shapes of a given object vary). Second, their shapes and colors were equally diagnostic of object identity (in nature, shape is more diagnostic of identity), and the colors across the set of objects evenly sampled color space (in nature, objects tend to be biased to warm colors). Third, the colors were calibrated, removing a confound of luminance contrast (in nature, yellow objects differ in both lightness and hue compared to red objects)^50,51^. Finally, the reward value of the objects was fixed, ensuring all the object concepts acquired the same subjective value (in nature, the subjective value of different objects varies for a given individual, and the same object can hold different subjective value across individuals).

The animals first completed alternative-forced choice trials in which the sample cue contains both the correct shape and color information of the object concept. The animals were rewarded with a drop of juice for touching the matching color or shape in the subsequent frame. We call these trials “short-term memory trials” because they can be successfully completed by storing in short-term memory the color and shape information of the cue sample in each trial without appeal to any long-term knowledge of the object concepts (**Fig. 1c**). The animals rapidly learned to identify the colors and shapes of the objects (**Fig. 1e**), attaining high performance (last 1000 trials, monkey W: 97.6%, [95% C.I.: 96.6%, 98.5%]; monkey J: 96.6%, [95.4%, 97.7%]), with no difference for color-matching versus shape-matching trials (see also **Fig. S1**).

After the animals reached >90% accuracy on short-term memory trials, they were challenged with sessions that required knowledge of the color-shape conjunctions of the objects (**Fig. 1d**). These trials use the same alternative-forced-choice delayed-match logic as short-term-memory trials but require knowledge of the long-term association of the object concepts. We therefore call these trials “long-term-memory trials”. Both animals performed above chance for the first 150 long-term-memory trials, suggesting that sustained engagement with short-term-memory trials had produced latent color-shape object concepts. To reinforce these conceptual representations, we interleaved sessions of short-term and long-term trials in many sessions per week over several years, mirroring the substantial experience children attain with stable concepts in school. Both animals reached high performance despite individual differences in the rate of learning (**Fig. 1f**; last 1000 trials, monkey W: 97.7% [95% C.I.: 96.8%, 98.5%]; monkey J: 87.3% [95% C.I.: 85.3%, 89.2%]), with no consistent difference between trial types at plateau.

### Brain areas that encode joint representations of color and shape

After the animals reached plateau performance on long-term-memory trials in their home cages, we had them perform long-term memory trials in the scanner (**Fig. 2a**). In each trial, a cue corresponding to the color (or shape) of an object concept was presented for 9 seconds, spatially jittered to mitigate retinal adaptation. The animal was then shown four colorless shape (or shapeless color) options and rewarded for making an eye movement to select the one linked by the object concept to the cue. The animals performed well (**Fig. S2**).

**Fig. 2:**
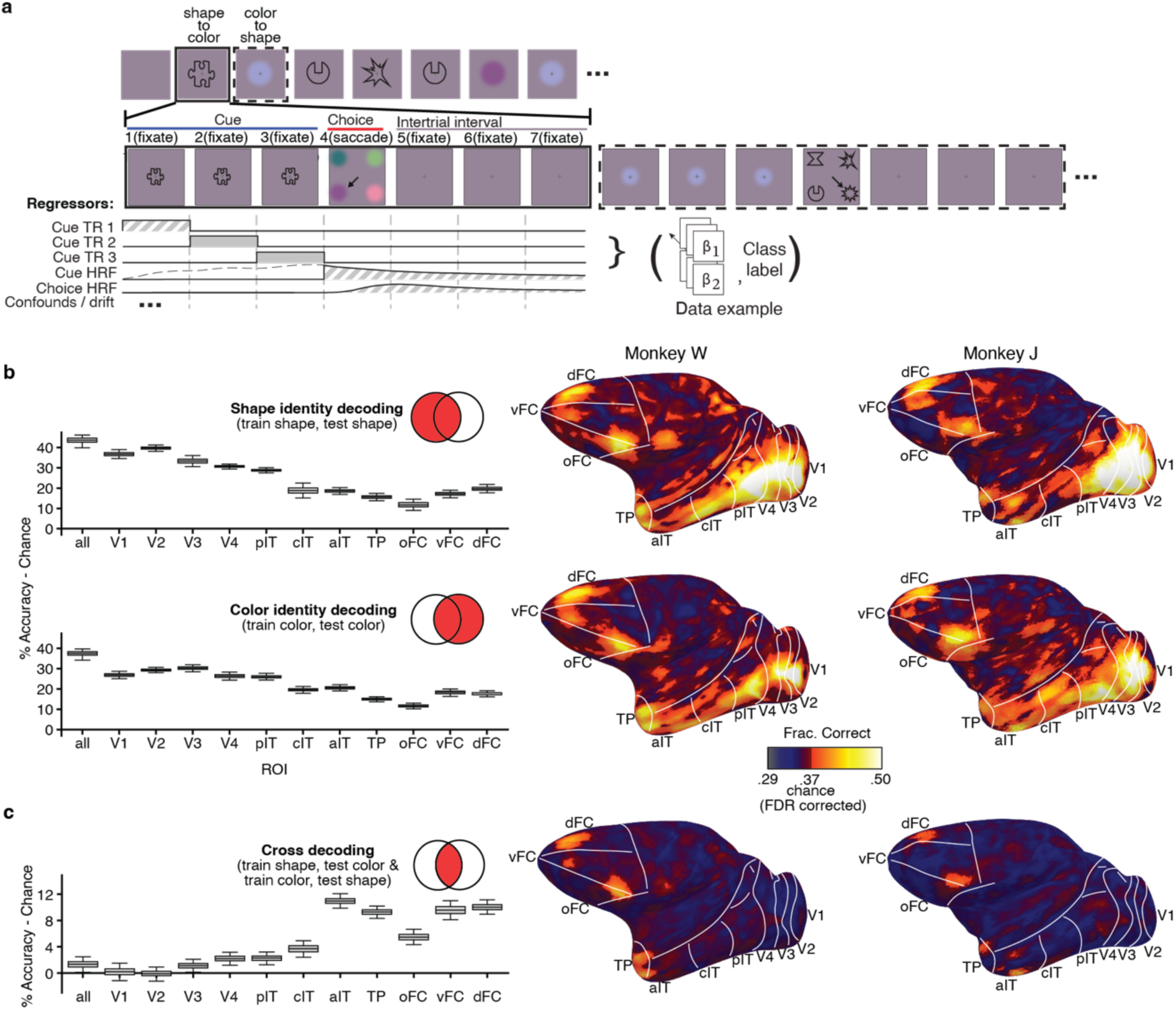
Representation of object concepts across macaque cerebral cortex. a, The sequence of trials in an example run conducted during fMRI (top), the frames during the TRs of two example trials in the run (solid box; shape trial, dotted box; color trial) (middle), and the regressors used in the general linear model (GLM) for the fMRI analysis (bottom). In each trial, the cue was presented for a total of 9 seconds (3 TRs), accommodating the hemodynamic delay of MION despite the lack of an HRF. The choice TR was followed by a 9-second inter-trial interval that allowed the fMRI signal to decay. Trials were in random sequence. Coefficients of cue TR2 and cue TR3 were used as inputs for the decoding analysis. **b,** Shape-identity decoding (top) and color-identity decoding (bottom), across the cortical surface of the left hemisphere of each monkey (right) and quantified in cortical parcels (left). Cortical parcels are: a grand parcel including all parcels (all); retinotopic cortical areas (V1 to V4); posterior inferior temporal cortex (pIT), central IT (cIT), anterior IT (aIT), temporal pole (TP), orbital frontal cortex (oFC), ventral frontal cortex (vFC), dorsal frontal cortex (dFC). Bars show percent accuracy minus chance; data is averaged across hemispheres and monkeys; error bars show 95% CI. Venn diagrams conceptualize the space of decodable outcomes. **c,** Cross-feature decoding (average of the two decoding problems, shape-to-color and color-to-shape); conventions as for panel (c). The color scalebar is the same for all surface maps. **Figs. S3 and S4** show results for all hemispheres, all decoding problems.

Our main goal was to determine the extent to which activity across the brain can support cross-feature decoding: when the animal is shown a colored blob and must hold in mind the shape of the associated object concept to successfully complete a trial, can we decode the identity of the unshown shape linked by the color-shape association, and if so where? Similarly, when the animal is shown a colorless shape and must hold in mind the color of the associated object concept to successfully complete a trial, can we decode the identity of the unshown color linked by the color-shape association, and if so where? To promote high detection power^52^, the paradigm was designed with key aspects of a block fMRI design. These include long cue presentations and long inter-trial intervals rather than relatively short cue presentations and short variable inter-trial intervals typical of event-related designs.

Importantly, we only analyzed data obtained during the cue TRs, not the choice TRs; and data were not analyzed with a hemodynamic response function (HRF) applied to the cue TRs. These analysis decisions eliminated any possibility that signals elicited by the choice were inadvertently assigned to the cue TR, which means that any successful cross-feature decoding must be ascribed to those signals “held in mind.” Throughout the main analyses, we also excluded the trials that the animals got incorrect or aborted, so we can be confident that during the cue presentation the animal was holding in mind the corresponding unshown feature of the object concept. On the basis of decoding work in humans^15,37^ and a quantitative comparison of data quality in monkeys and humans^53^, we set a target of ∼7000 trials per animal. We acquired 13,952 trials in the two animals over 48 scan sessions spanning nearly 100 hours of scan time.

Before evaluating cross-feature decoding, we first determined where across the brain neural measurements could decode feature identity (color or shape). Classifiers were trained on data acquired during the presentation of colorless shape cues and tested using independent data acquired during the presentation of colorless shape cues (evaluated using 8-fold cross-validation); and other classifiers were trained on data acquired during the presentation of shapeless colored blobs and tested using independent data acquired during presentation of the same shapeless colored blobs (again evaluated using 8-fold cross-validation).

The data were analyzed with a new searchlight decoder inspired by convolutional neural networks that we call the Layered Searchlight Decoding Model (LSDM) (**see Fig. 3** for validation of the LSDM compared to a standard searchlight). The LSDM is a form of multivoxel pattern analysis that discovers local structure and should be robust to noise. We configured the LSDM to provide probability estimates for each class on each trial within every 6x6x6 mm patch of the whole brain. The results were quantified in the 3-D volume. The surfaces (**Fig. 2b,c** right) show the decoding accuracy at each cortical location for the 3-D patch centered at that location. Note that area designations in surface projections are approximations because they require non-linear distortion, but they nonetheless provide landmarks to compare across data sets and analyses. To quantify the results, we aggregated searchlight patches comprising brain-atlas parcels in 3-D volume to generate box plots (see methods) (**Fig. 2b,c** left).

**Fig. 3.**
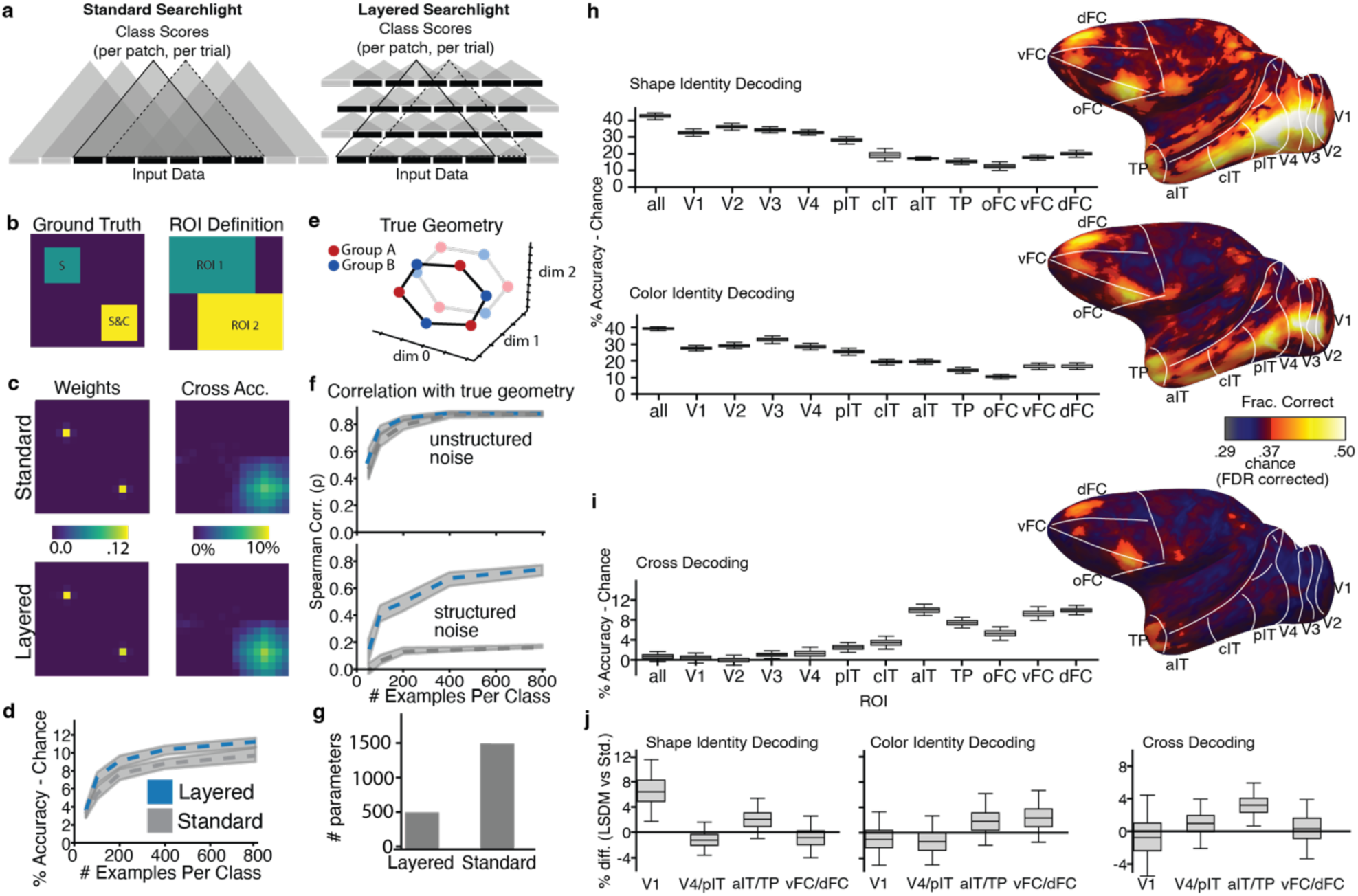
Comparison of Layered Searchlight Decoding Model (LSDM) and the standard searchlight using simulated fMRI data and real data. a, Schematic of the standard logistic searchlight architecture (left) and the LSDM (right). The first layer of the LSDM has filters (small triangles, bottom row), each with access to few voxels of input data. Subsequent layers are stacked so that the output filters comprising the searchlight patches (large triangles) each have access to a 6 x 6 x 6 mm cube of input data. **b,** For the simulation, the ground truth for cross decoding (left) and region of interest (ROI) definition. The upper-left (UL) region marked “S” encodes the identities of 12 “shapes”; and the lower-right (LR) region marked S&C encodes joint “color” and “shape” information. The two ROIs contain these ground-truth regions. **c,** The stacking weights learned by a decoder in the simulation (left) show the relative importance of each searchlight patch (sum to one over all patches). The above-chance cross-decoding accuracy (right) shows the results of train on C test on S. These results are illustrated on a plane through the middle of the input volume after fitting a standard searchlight (top) or the LSDM (bottom). Accuracy is higher for the LSDM. **d,** Cross-decoding performance of the LSDM vs standard searchlight model, for different simulated dataset sizes. The LSDM performs better. 95% confidence intervals are computed by bootstrapping model fits to independent data. **e,** Position of the different data classes on a low dimensional manifold of the input data for the simulation, matching the experimental paradigm (two circles of 6 colors at two different luminance levels). **f.** Correlation of dissimilarity matrices (RDMs) computed from the LSDM latent states, or via a standard representational similarity analysis (RSA) searchlight, with the ground truth geometry. The LSDM RDMs correlate better with the true geometry than the standard RDMs, especially in simulations that included class-dependent (structured) noise. **g,** Number of parameters per searchlight patch for the LSDM and standard searchlight in the simulation (number of searchlight patches is the same for both), matching the analysis of read data. **h,** Standard logistic searchlight identity decoding performance of real data. **i,** Standard searchlight cross-decoding performance of real data (compare with Figure 2). **j,** Difference in accuracy between the LSDM and standard search light on real data.

The results show strong shape-identity and color-identity decoding throughout retinotopic visual cortex (V1, V2, V3, V4), IT cortex (posterior, central, anterior parcels, pIT, cIT, aIT), the temporal pole (TP), and parts of frontal cortex (orbital, ventral, and dorsal PFC). Across parcels, decoding shape was slightly stronger than decoding color, but both shape-identity decoding and color-identity decoding peaked in V1 and V2 and declined progressively along the putative visual-processing hierarchy through the ventral visual pathway, rising again in frontal cortex.

**Fig. S3** shows surface maps for all hemispheres; **Fig. S4** shows box plots for all brain parcels including the hippocampus and subcortical structures. A direct comparison of the identity-decoding surface maps emphasizes the relatively greater decodability of shape in retinotopic cortex and reveals pockets of stronger decoding for color along the ventral surface of IT and PFC, consistent with prior work (Lafer-Sousa and Conway, 2013) (**Fig. S5**).

We then identified putative conceptual representations by training decoders with fMRI responses elicited by a given feature and testing with fMRI responses elicited by the other feature (the training and testing data sets are necessarily independent). Cross-feature decoding was tested in two ways: by training on shape data (and testing on color data) and by training on color data (and testing on shape data). **Fig. 2c** shows the average result. **Fig. S3b** shows the results for each separately and indicates the robustness of the results. The pattern of cross-feature decoding was strikingly different compared to identity-decoding. Cross-feature decoding was almost entirely absent within retinotopic visual cortex. Instead, it was evident in the anterior portion of aIT/TP and two locations of frontal cortex, one in dorsal prefrontal cortex (PFC), and one on the border of ventrolateral PFC and orbital PFC, peaking in area F5. There was significant but much weaker cross decoding in some other cortical and subcortical regions, notably the dorsal superior temporal sulcus, as well as in subcortical regions (**Fig. S4**). These results implicate aIT/TP and PFC as cortical candidates for the representation of color-shape concepts.

The decoding results in **Fig. 2** were generated with the LSDM, which we were inspired to develop because standard searchlights are computationally demanding. The LSDM and the standard searchlight, illustrated schematically in **Fig. 3a**, produce comparable patterns of results in both simulated data (**Fig. 3b-g**) and real data (**Fig. 3h-j**; See Methods describing the simulation to compare LSDM and standard searchlight methods). The LSDM achieved slightly higher accuracy at cross-decoding on both simulated (**Fig. 3d)** and real data (**Fig. 3j**), likely because it provides a form of regularization that makes it more robust to noise. The LSDM is also more memory efficient than the standard searchlight (**Fig. 3g**). The main conclusions from the standard searchlight are the same as from the LSDM: identity decoding was substantial across cortical parcels (**Fig. 3h**), while cross-decoding was substantial only in anterior IT, TP, and PFC; cross-decoding was weak or insignificant in retinotopic cortex (V1-V3), and significantly greater in aIT, TP and PFC compared to retinotopic areas (error bars are 95% CI) (**Fig. 3i**).

Note that it is impossible for signal from the monkey’s choice to inform the decoders because data acquired during the choice TR was not used in the analysis and no temporal kernel was used (see methods). Moreover, the decoding results also cannot be attributed to the animal’s choice in a prior trial, since the trial sequence was random. The decoding also cannot be explained by signals related to eye movements, since the correct choice on any given trial was at an unpredictable location. To confirm, we fit linear SVMs to eye-movement trajectories from each trial. Eye-movement trajectories could not reliably decode color-shape pair identity (8-fold cross-validation acc. – chance; W: .04 95% CI [0.01, .06], J: .02 95 % CI [-0.03, 0.07].

### Geometry of the color and shape representations across the cortex

The LSDM has an additional advantage over the standard searchlight: its results provide a latent space that helps uncover the geometry of the neural representation (**Fig. 3f**). We leveraged this advantage to determine the extent to which each parcel possesses a structured representation, with the goal of determining the contributions of different brain parcels to the conceptual representation. Colors are naturally ordered in a color circle: no explicit training is required to perceive similarity relationships among colors,^54^ and color-space geometry can be recovered from single-unit recordings in the V4 complex of monkeys^55^ and from fMRI data in a corresponding area in humans^56,57^. But many brain areas contain neurons that are responsive to color. Which areas possess a neural representation with a geometric structure corresponding to color space? To answer the question, we took advantage of the fact that in the LSDM, all data are projected to a low-dimensional manifold constructed such that the shape or color features of the concepts are optimally decodable. Signals (or noise) that carry no information about class identity relevant to decoding (i.e. within group and luminance-level class identity) will affect representational dissimilarity matrices computed directly on the input data, as is done in standard representational similarity analysis, but such signals will lie orthogonal to the latent space of the LSDM, which makes the latent space in the LSDM useful for performing a geometry analysis. This property of the LSDM analysis is similar to other neural population analyses where data are projected to task-relevant axes before analysis of representations or dynamics^58^.

We computed the matrix of pairwise Pearson dissimilarities (one minus the matrix of correlation coefficients) between stimulus classes (colors) from the final latent state of the LSDM generated for color-identity decoding. The dissimilarity matrix for a given visual area (**Fig. S6a**) was obtained as the weighted sum of the dissimilarity matrices of the LSDM patches comprising the area. Note that the projection LSDM is always fit on separate data than those used to compute the dissimilarity matrices. We ranked the pairwise dissimilarities among the six colors (equally spaced in perceptually uniform color space), allowing ties. We then correlated these two sets of rankings, obtaining the Spearman correlation (ρ) of the two dissimilarity matrices (**Fig. 4a**; **Fig. S6b** for light colors and dark colors separately). Because the correlation is computed on ranks of only 6 pairwise distances (averaged across four measures, light/dark for two monkeys), chance correlations can be non-trivial to compute. So, to estimate significance, we generated a null distribution for Spearman’s ρ by randomly permuting the model’s rank vector (preserving ties) 10,000 times; the gray bands in **Figs. 4a and S6b** show the 95% of this null distribution (2.5–97.5th percentiles). Correlations falling outside the band are significant at α=0.05. Boxplot error bars were generated via 8-fold cross validation. We plot the 2-dimensional Multi-Dimensional Scaling of the dissimilarity matrices to provide an illustration of the results (**Fig. 4a**, MDS plots labeled “color”). The correlations were significant in V3, V4, and IT, but not V1, consistent with the idea that perceptual color space is represented outside of V1^55,57^. The correlations were not significant in PFC.

**Fig. 4.**
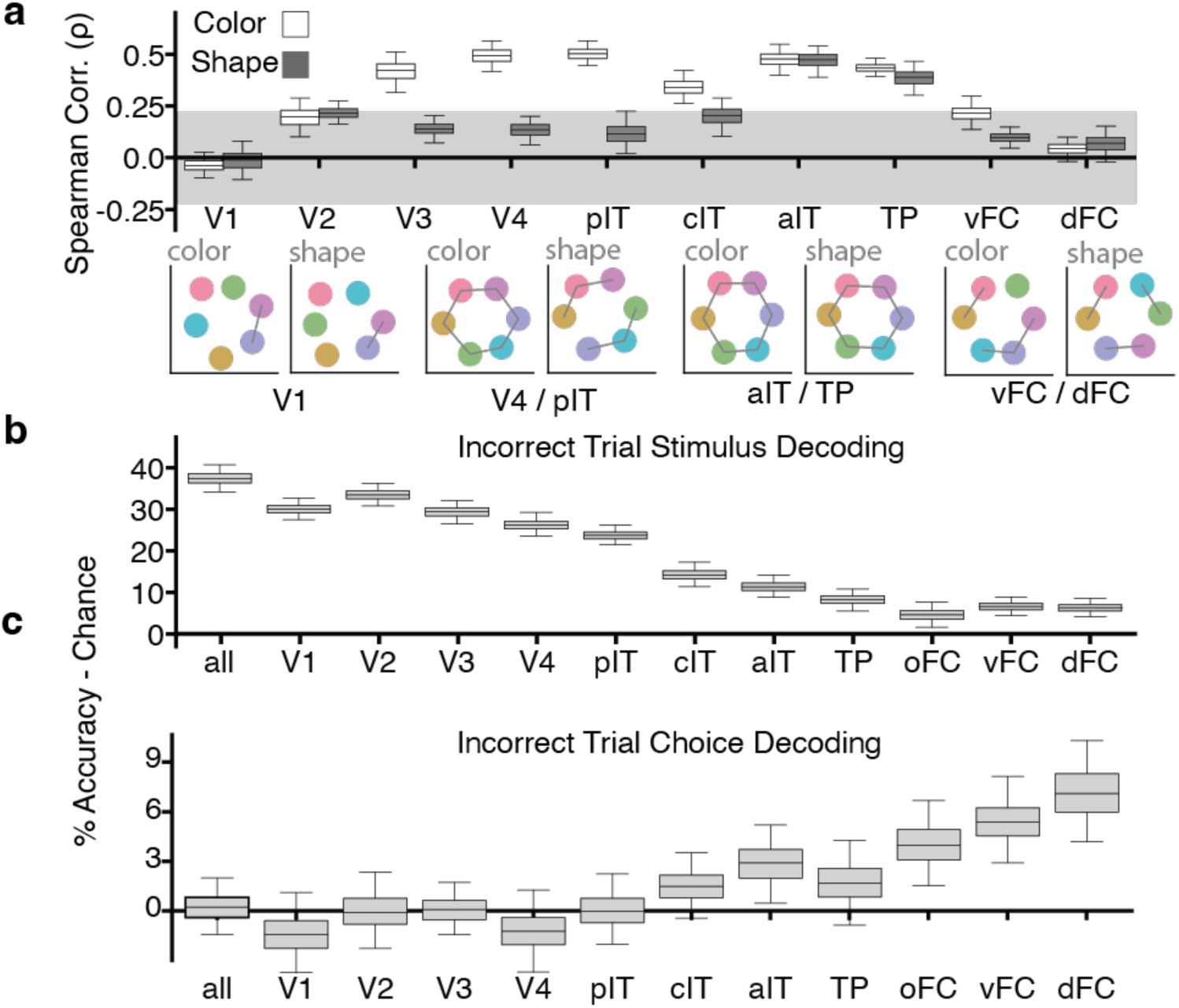
Structure of the neural representation of object concepts. a, Analysis of the pattern of fMRI responses collected while the animals performed long-term memory trials. Boxes show the Spearman correlation between the Pearson correlation distances in the LSDM latent space and the distances of colors in color space for standard cortical parcels (see Fig. 2 for key). Open boxes: correlation of the pattern of fMRI responses elicited by object colors with color-space geometry; filled boxes: correlation of the pattern of fMRI responses elicited by object shapes with color-space geometry. The shaded region shows the correlation level that could be achieved by chance (95% interval). Corresponding multi-dimensional scaling plots are shown below (grey lines connect neighboring items that are in the correct position predicted by color ordering of the geometry of color space). The geometry of color space given responses to object shapes is only complete in anterior inferior temporal cortex/temporal pole (aIT/TP). **b,** Accuracy of classifiers trained to decode the presented stimulus on error trials. **c,** Accuracy of classifiers trained to decode the choice the animal made on error trials; boxes show the average of color-to-shape and shape-to-color trials. Boxes in (b,c) show average results for both types of memory trials (color-to-shape, shape-to-color). Error bars show 95% C.I. by bootstrapping.

We next compared the representational geometry of responses to the colorless color-associated shapes with color space, to test for a structured conceptual representation, reasoning that a conceptual representation might use the scaffolding provided by the natural color geometry. We did this by computing the matrix of pairwise Pearson correlation distances between stimulus classes (shapes) in the final latent state of the LSDM generated for shape-identity decoding and ranking the pairwise Pearson correlation distance between exemplars in the latent model space. We then correlated this ranking with the ranking of the colors in color space (**Fig. 4a**, dark gray boxes). The only brain parcels to show a neural representational geometry matching color space were aIT and TP, not PFC. These results show that aIT/TP and PFC contribute differently to the conceptual representation.

A searchlight representational similarity analysis (RSA) without the LSDM’s latent space did not recover the representational geometries (**Fig. S6c**). In a simulation with known representational structure (**Fig. 3e**), the geometry is better recovered when using the LSDM’s latent space than when using standard searchlight RSA (**Fig. 3f),** especially when structured noise (noise dependent on class identity) is added (see “simulation to compare LSDM and standard searchlight” methods).

### Incorrect choices are represented in frontal cortex

In the final analysis of these data, we consider neural responses during the trials when the animal made a decisive but incorrect choice, “error trials”. As in all the analyses, we only included signal coincident with the cue TRs, i.e., before the monkey’s actual choice. We predicted that stimulus (cue) identity should be decodable from responses in visual cortex, since visual cortex encodes retinal data. But we predicted that the animals’ upcoming choices (even when incorrect), should be decodable from one or both of the areas that store a shared color-shape representation (aIT/TP, PFC). Whatever area(s) are recovered should identify it as the locus responsible for the final decision.

Consistent with our first prediction, the identity of the stimulus presented to the animal could be decoded with high success on error trials throughout visual cortex and IT (**Fig. 4b**). Meanwhile, the choices, though incorrect, were decoded best in the PFC, and most strongly in one of the two PFC foci, dPFC (**Fig. 4c**). These results further dissociate the cortical areas with strong cross-feature decoding. Taken together, the results support the conclusion that aIT/TP houses visual concept representations, while dorsal PFC reads out the concept representation to afford a final decision.

### Areas implicated in color perception and color-shape binding

Achromatic images of familiar color-diagnostic objects can be perceived as tinged with color by human subjects^59–62^. To the extent that monkeys are a model of the human case, we reasoned that brain areas involved in color perception might also show a bias for colorless color-associated shapes compared to colorless non-color-associated shapes. We tested this hypothesis in additional experiments in which animals were presented with blocks of colorless color-associated shapes and colorless non-color-associated shapes (**Fig. S7a**)—the animals were rewarded for passively fixating and not for making any judgements about the stimuli because we did not want them to update their conceptual representations. The animals had extensive experience with both colorless color-associated shapes and colorless non-color-associated shapes (**Fig. 1b**), so any differential response cannot be attributed to differences in familiarity. In all four hemispheres, the results showed a bias for color-associated shapes compared to non-color-associated shapes, and this response was observed only in extrastriate retinotopic cortex: V2 (color-associated shape selectivity = 0.057, 95% CI = [0.040, 0.075]), V3 (0.044, [0.028, 0.061]), and V4 (0.019, [0.004, 0.034]; **Fig. 5a**). It was not observed in V1 or in any region with strong cross-feature decoding. These results implicate extrastriate visual cortex in the conscious perception of color.

**Fig. 5.**
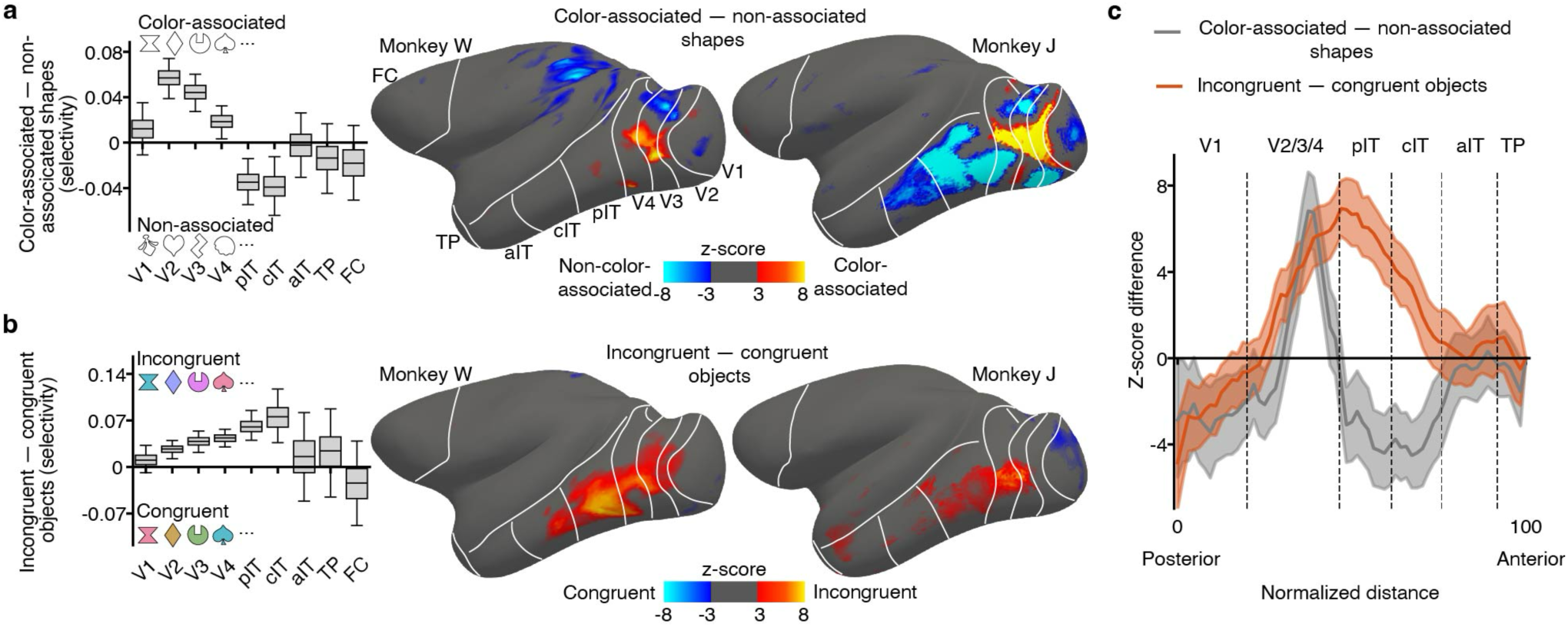
Areas implicated in color perception and color-shape binding. a, Differences in fMRI responses to colorless color-associated shapes versus colorless non-color-associated shapes across the cortical surface (right, Z-score contrast maps, bias for color-associated shapes in orange) and quantified in brain parcels (left, boxes show the difference between GLM beta weights divided by the sum of the beta weights for the two conditions on each run (analysis included 476 blocks per stimulus condition across 15 sessions for W, 244 blocks across 17 sessions for J; frontal cortex parcels anterior to premotor cortex are aggregated as FC). **b,** Differences in fMRI responses to colored shape stimuli that were incongruent versus congruent with the color-shape concepts (analysis included 112 blocks per stimulus condition across 4 sessions for W, 184 blocks across 5 sessions for J; other conventions as for panel a). **c,** Bias for incongruent stimuli (orange trace) and color-associated shapes (gray trace) on the same axes, computed along the posterior-anterior axis of the inflated surface. Shading shows 95% CI.

Prior work has hypothesized that color-biased regions of IT may be instrumental in object color knowledge retrieval^10,11,63–65^. We tested whether these regions show disproportionately greater responses to colorless color-associated shapes. The results do not provide compelling support for the hypothesis (**Fig. S8**): The bias for colorless color-associated shapes over colorless non-color-associated shapes did not appear to be restricted to color-biased regions, suggesting that computations that form color-shape associations depend on both color-biased and non-color-biased regions of IT.

Another long-standing question is how component features of objects are bound into a cohesive representation^66–68^. Such feature binding is not considered conceptual, but perceptual, because component features can be bound regardless of the conceptual content. The animals’ extensive experience afforded an opportunity to test for areas implicated in perceptual color-shape binding. In a final additional experiment, we compared fMRI responses to objects in which the colors and shapes were congruent versus incongruent with their learned object concepts (**Fig. S7b**). The power of this experiment derives from the perfect balance between conditions: the two conditions have the same colors and shapes, and the colors and shapes have been experienced with the same frequency and subjective value. We reasoned that any differential response must isolate regions that compute feature binding of color and shape. Again, the animals were rewarded for passively fixating to mitigate updating conceptual representations.

The results showed a bias for incongruent over congruent color-shape stimuli; this bias was only observed in V2, V3, and V4, and extending into IT, with a peak in posterior and central IT (TEO). It was not observed in V1 or any of the regions that showed strong cross-feature decoding (V2: incongruent object selectivity = 0.026, 95% CI = [0.012, 0.040]; V3: 0.039, [0.024, 0.054]; V4: 0.043, [0.029, 0.057]; posterior IT: 0.060, [0.038, 0.082]; central IT: 0.075 [0.035, 0.116]) (**Fig. 5b**). The regions implicated in color-shape binding are shifted anterior relative to the regions implicated in color perception (**Fig. 5c**).

To assess whether the results reflect a surprisal or novelty response, we analyzed separately the first and second halves of the data collected over a hundred blocks spanning several sessions. If the bias were caused by a difference in familiarity, then it should be stronger in the first half of the data. Contrary to this prediction, the bias for incongruent shapes was not only evident but also perhaps stronger in the second half of the data **(Fig. S9)**. Together with the results of the main experiment, the pattern of results suggests a visual-processing hierarchy, in which color perception is computed posterior to perceptual color-shape binding, and both perception and feature binding are computed posterior to color-shape concepts.

### Anatomical location of peak decoding and activation

So far, we have shown all the results on inflated cortical surfaces, these are useful for visualizing whole brain data, but can obscure exact boundaries and locations in cortex due to multiple stages of registration and averaging, and do not show subcortical areas. Therefore the main results from both experiments are shown on each monkeys high resolution anatomical volume (T1 scan) illustrated in **Figure 6** . High activation or cross-decoding accuracy regions are labelled with anatomical designation based on the NMT atlas ^69,70^. These labels are provided as a standard anatomical reference (areas in visual cortex shown elsewhere are derived from functional criteria as described in the methods). The locations of both activation (for the passive viewing experiments) and cross-decoding accuracy (for the active-task experiment) are remarkably consistent across monkeys. The bias for color-associated shapes is localized to extrastriate retinotopic cortex, with a peak in V2 for both monkeys. The bias for incongruent stimuli is also localized to extrastriate retinotopic cortex, extending into TEO/TEp, for both monkeys; in each individual, the peak color-associated shape bias is posterior to the peak incongruent stimuli bias. For cross-decoding, the peak of the vFC patch is localized to near the anterior border of premotor area F5 with area 12l (vlPFC) and the peak of the dFC patch is localized to area 9d/46d (dlPFC) in both monkeys. The peaks of the aIT and TP patches are localized to anterior TE and the granular temporal pole (specifically, TGsts) in both monkeys. There are also some individual differences. In monkey J the aIT cross-decoding patch is strongest in TEav, whereas in monkey W it is strongest in TEad and TEm. In monkey W there is a second peak of color-associated shape bias in V4, and a second peak of incongruency bias in TEpd. The incongruency bias is strongest for monkey W in TEO and strongest for monkey J in V4.

**Fig. 6:**
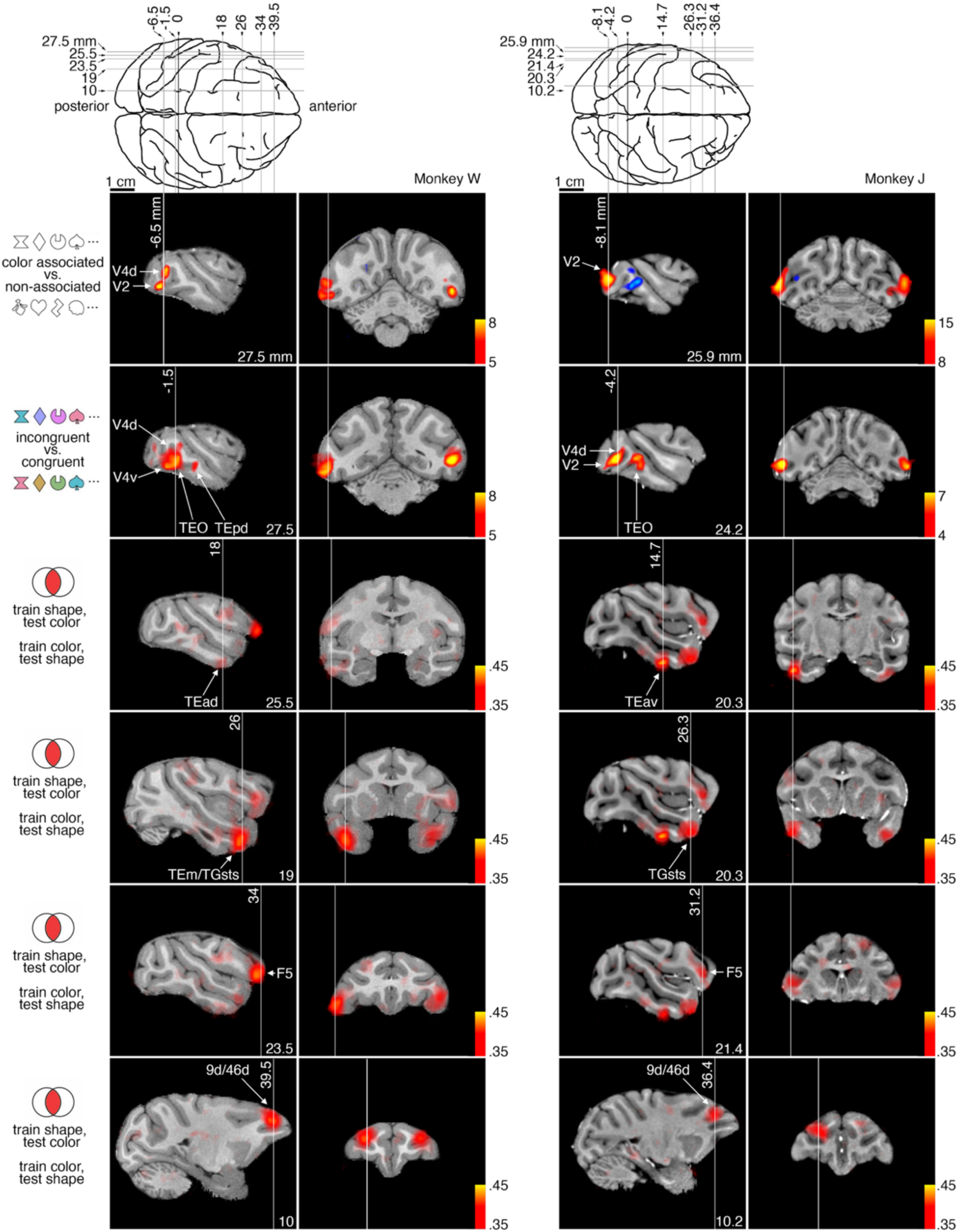
Anatomical location for regions of peak activation or decoding for three experiments. Top two rows: Color-associated versus non-color-associated shapes (see Fig. 5a) and color-shape objects that are incongruent versus congruent with learned color-shape concepts (see Fig. 5b); color scale is Z-score difference, threshold tailored so the underlying anatomy is not obscured. Bottom four rows: locations of peak cross-feature decoding, two within the temporal lobe and two within frontal cortex (see Fig. 2c); color scale shows decoding accuracy, chance = 1/3. Locations of slices are measured relative to ear-bar zero (coronal) and the midline (sagittal). Peak locations are labeled using the NMT v2 CHARM atlas.

## Discussion

The present work provides a missing whole-brain perspective on the representation of long-term, stable, visual paired associations that are the basis for concrete concepts. The results suggests a mechanism for how conceptual representations engage perceptual representations through selective feedback (**Fig. 7**), and provide a blueprint for how the brain constructs a memory of visual concepts (**Fig. 8**). The blueprint comprises a stage that encodes retinal images (V1), generates the perception of color and shape (V2, V3, V4), perceptually binds colors and shapes (culminating in TEO), houses conceptual color-shape associations (aIT/TP/FC), and registers decisions regarding conceptual content (dorsal PFC). The results provide a framework not only for understanding the normal mechanisms of visual cognition but also for interpreting synesthesia, mental imagery, and conscious visual experience.

**Fig. 7.**
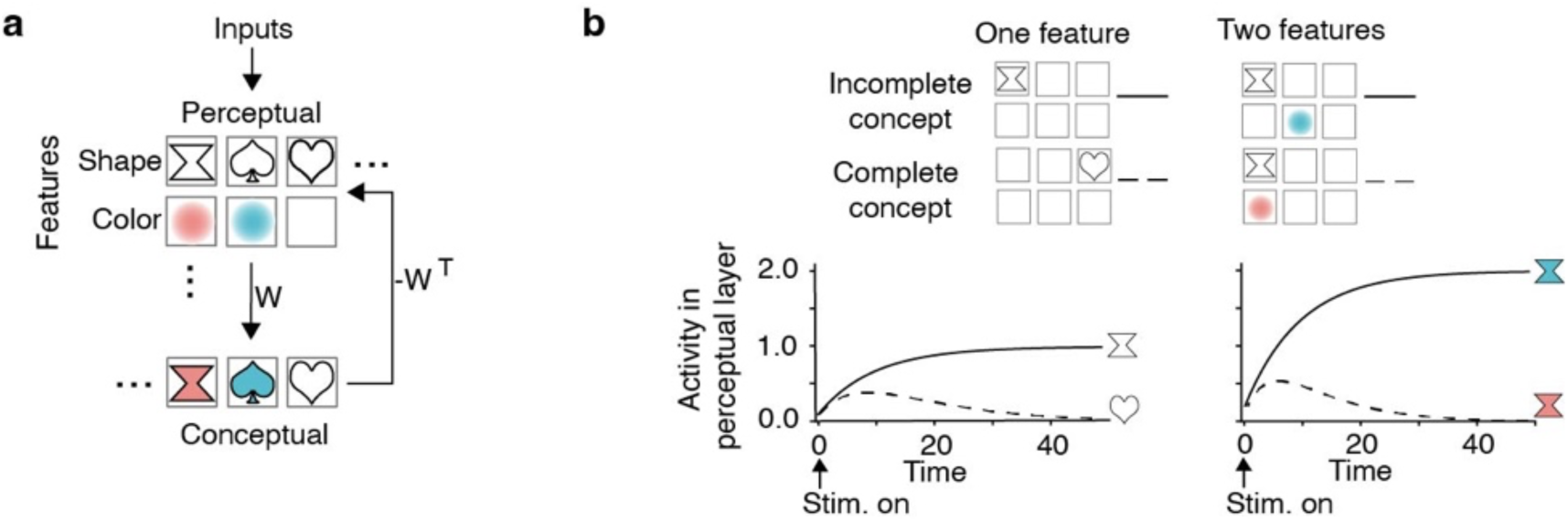
A simple model to interpret the results a, Schematic of the model, which has two layers, the first encodes separate representations for the perception of object features (shape, color), the second encodes a joint representation of the features as object concepts with weights, 𝑊. Feedback weights, 𝑊*, are the prediction of the perceptual layer activation given the activated conceptual representation. The perceptual layer computes the difference between the prediction and the received sensory data. **b,** In the model, colorless color-associated shapes and incongruent color-shape combinations constitute “incomplete” concepts—they only partially match a learned concept. For example, the hourglass outline is missing the color of the hourglass concept, and the teal-colored hourglass is the wrong color of the hourglass concept. Responses in the perceptual layer to complete concepts decrease rapidly, because the prediction matches the perceptual information; while responses in the perceptual layer to incomplete concepts are sustained, because the prediction does not match the perceptual information.

**Fig 8.**
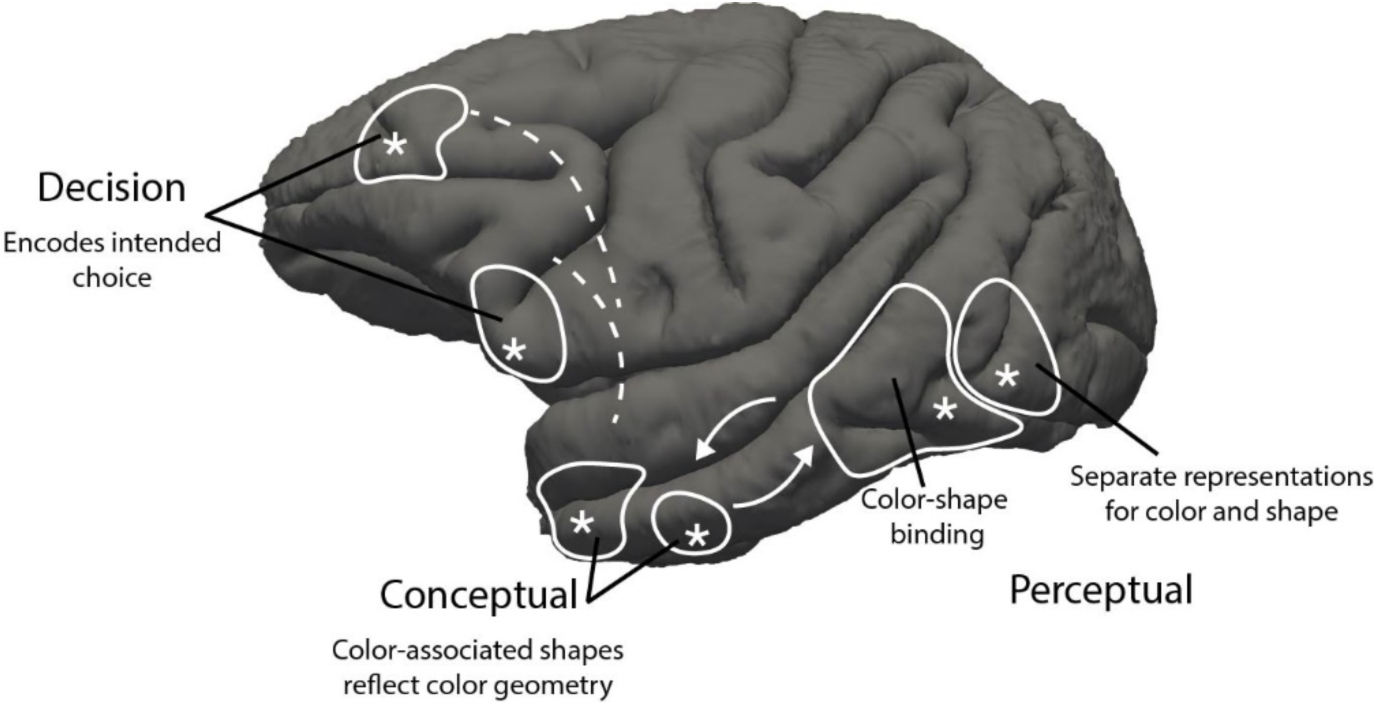
Hypothetical network for representing perception, concepts, and decisions about concrete visual objects. Contours show real brain activations in monkey W; asterisks show locations of peak activity or decoding (from in Fig. 6). Perceptual regions contain separate representations for color and shape (color-associated shape bias, Fig. 5a) that are then bound together (incongruent object bias, Fig. 5b). Conceptual areas house shared representations of color and shape (cross-decoding, Fig. 2c) wherein shapes are represented on a scaffolding of the perceptual color space (Fig. 4a). Decision-related areas contain shared representations of color and shape (cross-decoding, Fig. 2c) and represent the upcoming choice (incorrect decoding, Fig. 4c). Perceptual areas feed forward information about shown stimuli and receive feedback from conceptual areas (white arrows). Conceptual and decision-related areas communicate about unshown stimuli in service of goal-directed behaviors (white dotted lines).

The character, strength, and relevance of concepts vary among individuals, which has posed a challenge for understanding concept mechanisms. We overcame this challenge by raising macaque monkeys with a set of simple color-shape objects presented in self-initiated trials that taught the animals object concepts and provided them with behavioral enrichment. The paradigm offers precise knowledge of the color and shape content of the object concepts and controls the subjective value of the concepts, allowing us to determine from trials conducted during fMRI which brain areas “hold in mind” unshown concept features. We observed cross-feature decoding not in retinotopic cortex (V1-V4, or posterior IT), but in anterior IT, the temporal pole (TP), and dorsal and ventral foci in frontal cortex (FC). Cross-feature decoding was evident in both hemispheres, but slightly stronger in the left hemisphere. These results argue against the idea that retinotopic cortex is responsible for mental imagery, which is consistent with both the weak prediction performance of semantic features from retinotopic cortex^18^, and the observation that patterns of neural activity elicited from color words cannot decode color percepts^56^.

Among the three regions that showed strong cross-feature decoding, only aIT/TP showed a pattern of responses to concept shapes that was predicted by color space. This might seem at odds with prior observations showing a representational geometry in PFC reflecting the color circle^45^. But the two sets of results can be reconciled since PFC encodes task rules ^17^; in our study, the task rules made it advantageous to represent each color in isolation from the color wheel, while in the earlier study the task rules stipulated that color be represented in the color wheel.

The differences between aIT/TP and PFC uncovered by the geometry analysis (**Fig. 4**) show that these regions play different roles in the conceptual representation. But what does it mean for a conceptual representation to match color space? We think it suggests that the representation (in aIT/TP) is computed directly from an underlying perceptual representation, a conclusion that is supported by the anatomical position of aIT/TP at the culmination of the ventral stream where it receives feedforward input from the V4 Complex. Unlike a representational scheme comprising many separate dimensions, a scheme grounded in perceptual similarity could facilitate generalization to new concepts since it provides a representation of intermediate values. We therefore interpret the results as evidence that color space is used as a scaffolding for object concepts in aIT/TP, which could be exploited in new tasks.

It is presumably possible to represent color-shape concepts without either color or shape responses being predicted by color geometry. Such a representational scheme might make sense in tasks that treat each concept as a discrete entity (as in our task). Indeed, we believe the results indicate such a representation in frontal cortex, since the responses are not organized by color space.

Could a perceptual shape-space be used as a scaffolding for the conceptual representations? We could address this question if we knew the ground truth shape-space geometry, but this information is unknown. The shape-space geometry is likely high dimensional. Our working hypothesis is that the brain exploits color to represent object concepts, not only in the present tasks but also in natural behavior, because color has a relatively low-dimensional perceptual geometry. If true, this hypothesis could explain why so many conceptual associations are forged with color, as illustrated by the preponderance of cases of synesthesia ^71^, and the fact that industry logos often exploit color, color is often used in metaphor (e.g., “green with envy”, “sound color”), and in personal, professional, and political identities. These observations imply that color is generally useful as a scaffolding for concepts, which would also predict that when conceptual representations consist of many features defined by different feature spaces, the conceptual representation will be governed by color. We hope to test this prediction in future work.

In additional experiments, we determined which areas distinguish colorless color-associated shapes from colorless non-color-associated shapes, and which brain areas distinguish color-shape combinations that are congruent versus incongruent with the learned color-shape concepts: retinotopic extrastriate visual cortex (V2, V3, V4) distinguished color-associated from non-color-associated shapes, while brain regions extending anteriorly into IT reflected knowledge of color-shape combinations that were congruent with object concepts (**Fig. 5**). Do these results reflect a surprisal or novelty signal? We do not think so because they do not involve brain regions typically engaged by novel or surprising stimuli^72^ and do not decay with increasing familiarity (**Fig. S9**).

Instead, we believe these results reflect feedback from areas that house color-shape concepts, as illustrated in Figure 7—these regions do not themselves appear to house the concept representations because they did not support cross-feature decoding. This interpretation assumes that extrastriate retinotopic cortex would show cross-feature decoding with fMRI if such representations existed. This assumption might be invalid if, for example, the cross-feature representations were not appropriately organized to be detected by fMRI. The notion of separable brain areas for perception and cognition of color is predicted by studies of brain damaged patients^73^ and imaging of blind and sighted people^74^.

Selective feedback could take many forms. For example, it could reflect feature attention^75^, an error signal between expectation and perception^76^, or a prediction about what is expected^77^. Our data implicate some form of feedback but do not distinguish these mechanistic accounts. But to make our explanatory framework concrete, we illustrate it with a two-layer model (**Fig. 7**). In the model, colorless color-associated shapes and incongruent color-shape combinations constitute “incomplete” concepts—they only partially match a learned concept. In the model, a learned concept (an “expectation”) is directly compared to an incoming perceptual signal, and the difference is propagated through the network. One way of thinking about the model’s operation is that the conceptual layer sends a feedback “request” for data specifically about the missing perceptual information when confronted with an incomplete concept.

For a colorless shape that is typically associated with a color, the feedback requests information specifically about the perception of color, since color is the missing feature, while for a colored shape consisting of a color-shape combination incongruent with an object concept, the feedback requests information specifically about the perceptual binding of colors and shapes. Accordingly, the fMRI data implicate extrastriate retinotopic cortex as the locus for conscious color perception and an overlapping region extending anteriorly into TEO as the locus of conscious perceptual color-shape binding.

This account is consistent with theories of cortical function^76^, and is appealing for several reasons. First, it accommodates the fact that observers can consciously perceive colors and shapes of objects independently—these percepts provide separable information such as object identity (shape) versus object state (color). Second, the framework shows how the brain could learn to link features (paired associations) that are not reflexively bound in perception—this provides a visual system that can simultaneously see and think about what is seen. Third, it helps explain how feedforward and feedback signals could be engaged during normal behavior. Our working hypothesis is that during undirected vision, retinal data is encoded by feedforward signals to generate perception; the percepts are reflexively compared against the vast bank of concepts stored in aIT/TP, which provides selective feedback to perceptual stages whose activity is needed to connect what is seen with what might be desired; and finally, PFC dictates the task rules and reads out perceptual decisions (**Fig. 8**). This account also informs longstanding discussions about whether the sensitivity of many V1 neurons to spatial chromatic structure reflects mechanisms for perceptual color-shape binding^78–81^—the present results suggest it may not.

## Materials and methods

Preliminary presentations of the present work were given earlier^82,83^.

### Longitudinal experience with color-shape concepts

#### Overview and stimulus design

Two juvenile male macaque monkeys (J and W, born 07-20-2014 and 08-07-2014) were provided color-calibrated touchscreen tablets in their home cage. The touchscreens were introduced on 01-27-2016. The touchscreens were connected to a fluid reward delivery system that provided drops of juice from a spout in front of the tablet. The animals used the touch screens in self-initiated trials to learn a set of “objects” consisting of combinations of colors and shapes, or just colorless shapes (**Fig. 1b**; two additional objects, light and dark gray, were also included but not analyzed). Such simple 2-dimensional stimuli have been called “concept primitives”^49^; we refer to their neural representation as “object concepts.” We recognize that these stimuli do not constitute “objects” in the fullest sense of the concept of objects—for example, the stimuli are not three dimensional, they are only “handled” using a touch screen, and the degree to which they are experienced in many contexts is reduced compared to the natural experience of objects. Nonetheless, the animals saw the stimuli from multiple angles in their home cage, their engagement was self-initiated throughout the day (i.e. in many behavioral contexts), and they received reward in direct response to touching the objects.

One challenge in studying object concepts is their individual differences in character, strength, and behavioral relevance. For example, a person’s concept of “banana” might be a solitary piece of ripe fruit in profile, a half-pealed piece of fruit ready for consumption, or a bunch of fruit hanging from a tree. In addition, people have different ideas about the color of the banana concept—is it the most frequently encountered ripe banana, the tastiest banana, or focal yellow? Moreover, people do not like eating bananas to the same extent, which means the concept holds a different subjective value for different people. The paradigm overcomes this challenge by controlling the visual features and reward value underlying the object concepts. Humans could easily learn the tasks of the paradigm, but since our objective is to understand the neural representation for stable long-term concepts, it would be necessary for humans to engage the tasks for years and to derive substantial reward from them, which was not feasible.

The paradigm controlled six dimensions:

1. The colors evenly sampled the hue circle (CIELUV), which controls for a bias in the color statistics of natural and artificial objects ^51^. This control enabled a test of the structure of the neural representation (see **Fig. 4**).
2. The colors controlled for luminance contrast. Half of the colored objects were a fixed higher luminance contrast compared to the background, and half of them were a fixed lower luminance contrast than the background. This removes a confound in natural objects where hue and luminance contrast are often correlated. For example, yellow objects are typically a different hue and a different luminance contrast than red objects.
3. The shape features were restricted to simple feature elements drawn with lines, which controls object complexity. The object shapes were drawn by us, with the objective of having shapes that were well discriminable from each other and matched in visual interest. We realize these are somewhat arbitrary criteria, so in initial pilot experiments we measured fMRI responses in other animals to eight candidate sets of shapes, and we selected for the present work the two sets that elicited comparable, strong, responses.
4. The feature contingency of the objects was fixed, so the paradigm enables the acquisition of stable concepts akin to those learned by children in school. Such concepts can be accessed by independent features, which resolves a confound that has been a challenge in non-linguistic work on concepts in humans^9,21^.
5. Both color and shape were well above discrimination threshold and were equally diagnostic of object identity. Since the colors and shapes are precisely defined, there is no uncertainty about the ground truth for the corresponding mental representation.
6. The subjective value of the different concepts was controlled since each concept was linked to the same amount of the same juice. This control is difficult with natural objects because the reward value of objects differs among people.

There were two phases of behavioral testing in the present work. In phase I, trials could be completed solely using short-term visual memory: animals were rewarded for directly matching the identity of the color or shape of the objects in an alternative-forced-choice paradigm (**Fig. 1c**). Animals initiated a trial by touching a textured “button” on the tablet that triggered the immediate presentation of one of the objects at the center of the tablet. After 500 ms, the object disappeared, and choice options were presented. If the object was colored, the choices could be either colorless shapes matching the shape of the object or shapeless colored blobs matching the color of the object; if the object was one of the colorless objects, the choices were drawn from the set of colorless objects. The three trial types were randomly interleaved. The paradigm initially provided two choices (2-Alternative-Forced-Choice, 2AFC). The animals rapidly reached high performance. To increase task difficulty, we adapted the paradigm to four choices. Animals performed an average of 844 trials per day and rapidly reached high accuracy. A detailed assessment of the behavioral performance will be provided in a separate report.

The colors and shapes of the objects were designed to be balanced in supra-threshold discrimination. We tested this by measuring behavioral responses to a subset of 2AFC short-term memory trials in which the animals directly matched colorless shapes to colorless shapes, and shapeless colored blobs to shapeless colored blobs. At plateau performance, there was no difference in performance between matching color and matching shape trials (**Fig. S1**, % correct color-to-color, % correct shape-to-shape, [95% CI], for the last 1000 trials: W, 93.3% [91.7, 94.9], 92.4% [90.6, 94.0]; J, 91.2% [89.6, 92.8], 88.8% [86.9, 90.8].

In phase II, successful trial completion required long-term memory of the color-shape associations of the objects (**Fig. 1d**). Phase II trials were routinely introduced when the animals had reached at least 90% accuracy on 4AFC trials of phase I, i.e. when they had sufficient exposure to have mastered reporting the colors and shapes of the objects and, we reasoned, had acquired latent color-shape concepts of the objects. There were two trial types in phase II: one in which a colorless shape matching the shape of one of the objects was presented as a cue, followed by match options consisting of shapeless colored blobs (“shape-to-color” trials); and one in which a shapeless colored blob matching the color of one of the objects was presented as a cue, followed by match options consisting of the colorless shapes (“color-to-shape” trials). The animals were rewarded for making matches determined by the color-shape associations forged by familiarity with the objects.

Although one might assume that any latent color-shape object concepts were the result of learning phase I, we considered the possibility that in setting up the experiment, the color we assigned to each shape reflected an unconscious bias we had about what color the shapes should have – perhaps the colors we pick for each shape are influenced by unrecognized statistical associations in the natural world (perhaps puzzle pieces are dark purple and multi-pointed stars light are lavender). So, out of an abundance of caution, we collected two sessions of long-term memory trials prior to beginning short-term memory trials, i.e. before the animals had seen the colored shapes. The animals were, in fact, at chance on these trials (% correct shape-to-color, % correct color-to-shape, [95% CI], number of trials: W, 49.9% [41.5, 57.7] n = 142, 54.8% [42.6, 66.2] n = 68; J, 57.9% [46.8, 68.3] n = 79, 39.2% [26.3, 55.3] n = 38, chance is 50%), which supports the conclusion that the color-shape associations of the colored objects was arbitrary, as intended.

Error bars in **Fig. 1e** and **Fig. 1f** show 95% C.I. computed by bootstrapping the data binned by each data point.

#### Husbandry and fMRI procedures

The animals were pair-housed with standard 12:12 light:dark cycles and ample enrichment provided by toys, fruits, radio and television. During behavioral testing days, the animals were separated for a few hours to provide each animal with its own touch-screen tablet testing box. The animals received most of their fluids through performance of tasks on the touchscreen tablets. Their fluid intake was closely monitored to ensure they received at least the minimum requirements to remain healthy; the minimal requirements were established in consultation with veterinary staff and by measuring water consumption on days when the animals were given free access to water. The fluid consumed during behavioral-testing days regularly exceeded minimal requirements; indeed, the animals routinely continued performing trials long after the fluid was depleted from the reward system, suggesting that the animals found the task inherently rewarding.

The procedures we used for conducting fMRI with macaque monkeys have been described previously^63^. Briefly, the animals were prepared for imaging and trained using positive reinforcement to enter a custom-designed plexiglass chair that fit in the bore of a standard Siemens 3T MRI machine identical to ones used for human imaging. The animals were oriented in a sphinx position with their eyes facing a screen at the end of the bore, send-receive coils were placed around the head, and a tube connected to a reservoir of juice was positioned near the animal’s mouth to deliver fluid reward during scanning.

### Tasks in the fMRI experiments

#### Active task: Engagement of the learned color-shape object concepts

The experiments using this task provide the data for the main decoding analysis (**Figs. 2-4, 6**). We required the animals to actively engage the object concepts (instead of a passive fixation), for two reasons: first, to supply behavioral control to ensure that the animals were, in fact, holding in mind the concept features during imaging; second; because we reasoned that active engagement would increase the fMRI signals associated with the concept representations.

The active task consisted of long-term memory trials adapted in two ways for the scanner: first, the animals could use an eye movement rather than a finger to select their choice; second, the trials had longer epochs of cue presentation and inter-trial intervals to accommodate the hemodynamic delay of the fMRI signal (described in detail below). The animals were trained with positive reinforcement to fixate a spot on a screen at the end of the scanner bore. The animals generalized performing the trials from their home cage (with touchscreen) to the MRI machine (with eye movement) after a few training sessions. Note that all fMRI experiments were performed long after the animals had acquired expert knowledge of the object concepts through the touchscreens in their home cage. **Fig. S2b** shows the accuracy of completed (non-aborted) trials, for each cue, for each monkey; **Fig. S2c** shows the proportion of aborted trials. We attribute these aborted trials to the fact that trials in the scanner were not self-initiated. The total proportion of aborted trials were 5.7% (monkey W) and 4.6% (monkey J).

As with the long-term memory trials completed using touch screens, there could be two trial types. In the “color-to-shape” trial type, the animals were shown for three TRs (3 sec/TR) a colored blob cue selected from the set of colors of the colored-shape object concepts; on the fourth TR they were shown four colorless shapes and rewarded for making an eye movement to the option that matched the object concept. In the “shape-to-color” trial types, the cue was a colorless shape selected from the set of shapes of the colored-shape object concepts, and the match options were colored blobs.

In all trials, the TR containing the choices was followed by three TRs of gray, so the inter-trial interval was 9 seconds (**Fig. 2a**). Including the inter-trial interval, each trial was therefore 7TRs, or 21 seconds. The cue was presented at the center of gaze, spanning ∼3 degrees of visual angle (diameter), but with spatial jitter of ∼ 1 degree of visual angle across time to mitigate adaptation. The display provided the animals with a fixation cross throughout the cue TRs. The fixation cross disappeared as soon as the match options appeared. The match options were presented at the four corners of a virtual square with 3.2 degrees of space between edges of neighboring options, centered on the location of the fixation spot. The location of the correct choice was random on each trial. A small juice drop (10 ms open time on the solenoid controlling flow) was given at random times throughout the trial to encourage the animals to stay on task, and a substantially larger juice drop (150 ms open time) was given for making a correct choice. The reward value for successful completion was the same for all trials. Fixation was monitored with an infrared camera (ISCAN, 30 frames/second), and trials were aborted if the animal’s eyes deviated during the cue TRs by more than 1 degree from the fixation spot.

Based on prior work, we estimated the amount of data needed to do cross decoding. One study that aimed to decode the identity of objects using color and shape attained significance with a total of 528 trials across four trial types and 11 human participants^15^ (132 repeats per trial type). A quantitative comparison of fMRI signals in humans and macaque monkeys shows that one needs perhaps ∼8x or more data in monkeys, depending on the individual animal, to achieve the comparable data reliability^53^. These considerations gave a ballpark estimate of 1,000 repeats of each trial (∼8x132). But since we intended to analyze data separately for each animal, we assumed the ballpark estimate to be lower. Knowing how much lower requires information about the variability in the neural representation patterns across individuals, which could be caused by both genuine individual differences in patterns of brain activity and differences in data reliability. This information is unknown, so we made a best guess, setting a target of 250 repeats per trial, per animal.

We define a trial pair as the shape-to-color trial and the color-to-shape trial that relate to the same concept. Our experiment included 14 trial pairs, 12 of which involved colorful shapes and 2 of which were gray shapes (one was lighter than the background, the other was darker), for a total of 28 trials. Data from the 2 trial pairs involving gray concepts were not analyzed for two reasons. First, gray stimuli occupy a fundamentally different position in color space than the other colors; and second, one animal refused to engage with the gray concepts in the scanner even though they had happily learned the gray concepts in their home cage. Across all trial pairs, the estimated target number of trials was: 250*28 = 7000 trial presentations per animal, which is ∼80 hours of scan time not including set up, dummy scans, and aborted trials (7000trials/animal * 2animals * 7TR/trial * 3sec/TR * 1hour/3600sec).

To generate the sequence of trials for each run in the scanner, we aimed to control for both the time during the run when a given trial was presented and the number of repeats of each trial. We could not accommodate all trials in a single run, because of limitations of the scanner (and the monkeys). We initially adopted the following procedure, which we used for 10 sessions in one animal and 4 sessions in the other animal until it became clear that it would not provide the same number of repeats per trial. For the first run, we randomly selected four trial pairs and used a second-order deBruijn sequence to counterbalance trial order ensuring each cue pair (in order) appeared once^84^. The deBruijn sequence produced four runs each consisting of 17 trials, with the first trial of each block being a repeat of the last from the previous block to maintain counterbalancing. All runs included three additional TRs at the beginning and four additional TRs at the end, during which we showed the animal the gray screen. In each trial, the four choice options were determined by the four concepts to which the trial pairs corresponded. When the four runs were completed, we generated a new series of runs, by again randomly selecting four trial pairs from the set of unique trials and repeating the procedure with deBruijn counter balancing. In each session, we aimed to complete as many runs as the animal would perform.

After 10 sessions in one animal (W), we had obtained data on an average of 121 repeats of each trial [s.d. 44, range 37 – 217; 3383 trial presentations]; after 4 sessions in the other animal (J), we had obtained data on an average of 33 repeats of each trial [s.d. 15, range 8 – 61; 924 trial presentations]. To enforce a more consistent number of repeats of each trial, we switched the procedure for selecting trials on each run and used longer runs. We divided the trials into two groups (A and B), such that each group had an equal number of light and dark colors and evenly sampled the hue circle (**Fig. S2a**).

In monkey W, we collected 6 sessions of group A and 6 sessions of group B. Two sessions of set A and one session of set B were excluded from analysis because of massive artifacts caused by the scanner overheating. In monkey J, we collected 19 sessions of group A, and 6 sessions of set B. We collected more sessions of one of the sets in monkey Je because this animal showed poorer performance.

We obtained the following data:

Monkey W, group A: 152 repeats of each trial [s.d 6, range 146-166] Monkey W, group B: 131 repeats of each trial [s.d 4, range 122-138] Monkey J, group A: 115 repeats of each trial [s.d 4, range 107-123] Monkey J, group B: 272 repeats of each trial [s.d 21, range 240-306] Note that the slight variability in trial repeats arises because of the cyclic wraparound introduced by deBruijn counterbalancing and the fact that within a given session, we could not obtain complete sets of runs due to variability in animal performance.

Across all sessions, we obtained for all trials an average of 263 trial repeats in monkey W, and 227 trial repeats in monkey J. As described below, the variability in repeat numbers for each trial was addressed in the analysis by oversampling (after train / test splits) during cross-validation and by dynamically adjusting the weight given to each class based on its difficulty, as done by others^85^.

As described above, by chance some trials were slightly more frequently presented to the animals in the scanner. We were curious if the animals performed better on these more frequent trials, as one might expect if they were still learning about the objects—this concern is raised because monkeys are notoriously poor at generalizing, so it is conceivable that they were confronting the trials in the scanner as if they had had limited experience with the objects.

Contrary to this possibility, the accuracy of completed trials as a function of the number of trial repeats shows no correlation in one animal (r=0.19, p=0.37) and a weak *negative* correlation in the other animal (r=-0.4, p=0.05) (**Fig. S2d**). Together with the excellent accuracy on the trials in the scanner, we interpret this as evidence that the animals had generalized the concept knowledge from home-cage training. Our interpretation of the slightly negative correlation in one animal is that it was so familiar with the concepts that it was slightly above chance in keeping track of the concept frequency and was expecting to see a consistent number of repeats of each trial as they had in their home cage.

#### Passive-viewing task 1: fMRI signals related to the color association of colorless color-associated shapes

Much of human visual behavior does not involve intentional retrieval of concepts but rather the feedforward computation of visual stimuli coupled with un-directed processing that relates these signals to the vast bank of concepts stored in the mind. We sought to investigate this processing, guided by research showing humans sometimes perceive slight tints to achromatic color-diagnostic objects ^59–62^. To investigate neural activity related to the color-shape conceptual primitives that the animals knew well, we conducted fMRI scanning of the two animals while they passively viewed blocks of colorless color-associated shapes, colorless non-color-associated shapes, colored shapes, shapeless colored blobs, and full-field gray. **Fig. S7a** shows the fMRI paradigm. The animals were rewarded simply for maintaining fixation, ensuring passive engagement and mitigating the risk that the animals updated their repertoire of object concepts (if the animals were rewarded for making judgements about the stimuli, then they would begin to form new conceptual representations of them). All block types appeared in every run, and the order of block types was counterbalanced across runs using deBruijn sequencing^84^. Our analysis focused specifically on blocks showing colorless color-associated shapes and blocks showing colorless non-color-associated shapes, since these blocks comprise the same visual features (familiar colorless shapes), but differ in terms of the animals’ color association.

Within each block, all 14 stimuli of the corresponding stimulus condition were shown in a randomized order. Each stimulus was presented for one TR (3 sec), at the center of gaze. As in the active task, the stimuli spanned ∼3 degrees of visual angle, with spatial jitter of ∼ 1 degree of visual angle across time to mitigate adaptation. The monkeys received juice rewards for maintaining fixation on a cross at the center of the display.

In the first sessions for each animal, we used 20 blocks per run and collected 6 sessions in monkey W and 1 session in monkey Je. These runs were long, exhausting the scanner and the animals, so for subsequent sessions we reduced the run length to 10 blocks per run. The sequence of blocks for three runs is shown in **Fig. S7a** . We collected an additional 9 sessions in monkey W and 16 sessions in monkey Je. Note that the counterbalancing ensures that, regardless of run length (as long as at least one 5-block sequence containing all block types is completed), across the set of runs collected, every possible ordering of two block types appeared an equal number of times. We excluded any run in which fixation dropped below 80% on any stimulus block. This exclusion criteria yielded a total of 476 blocks per condition (across 15 sessions) for monkey W and 244 blocks per condition (across 17 sessions) for monkey Je.

#### Passive-viewing task 2: fMRI signals related to perceptual color-shape binding

Guided by the same logic as the first passive-viewing task, we sought to determine brain areas that responded selectively to the perceptual binding of colors and shapes. [Note that this experiment depends implicitly on a distinction between “perceptual binding” and “conceptual binding”: the former involves the perception of the conjunction of colors and shapes independent of whether the colored shape has been seen before; the latter requires memory.] After all active choice and passive-viewing task experiments were completed, we scanned the monkeys while they viewed alternating blocks of “congruent” objects and “incongruent” objects, interleaved with full-field gray blocks (**Fig. S7b**). Congruent objects were simply the 12 color-shape object concepts that animals knew well: the color and shape of each stimulus was congruent with the color-shape pairing of one of the concepts. Incongruent objects were generated by swapping object colors across CIELUV, separately for each luminance level. So, the light yellow of the diamond was swapped with the light lavender of the multi-point star, resulting in an incongruency between the color and the shape as compared with the concepts that the animals knew well. This one-to-one swapping of colors within luminance level ensured that each shape and color was represented only once in both the congruent and incongruent set, and that the two sets differed only in the shape-color congruency (not luminance).

Each run consisted of 16 blocks; all 12 objects were presented in each block in random order. This paradigm results in a perfectly balanced design in which every congruent block contained precisely the same shapes and colors as every incongruent block, and that the colors and shapes were all equally familiar to the animals. Such control is difficult to achieve in experiments with human subjects^86,87^. The objects were presented at the center of gaze for one TR (3 sec), spanning ∼3 degrees of visual angle, with spatial jitter of ∼1 degree of visual angle across time to mitigate retinal adaptation. Over the course of each run, subjects received juice reward for maintaining fixation on a fixation cross at the center of the display. Runs with lower than 80% fixation on congruent or incongruent stimulus blocks were excluded from analysis. We included a total of 112 blocks per condition (across 4 sessions) for monkey W and 184 blocks per condition (across 5 sessions) for monkey Je.

### Localization of retinotopic areas and functional regions

We performed three additional fMRI experiments to identify functional divisions of visual cortex in the two animals.

First, to define boundaries between retinotopic areas, we conducted a standard meridian mapping paradigm consisting of alternating blocks of horizontal and vertical checkerboard wedges^63^.

These stimuli elicit strong responses along the horizontal and vertical meridian representations in visual cortex, and their contrast reveals retinotopic area boundaries. The paradigm was identical to the one we used previously^63^. We obtained the following data: W, 18 runs, 1 session; Je, 17 runs, 2 sessions.

Second, we conducted an experiment to identify the eccentricity representations (foveal versus periphery) in the two animals, again using a standard paradigm^63^. Black and white checkerboard rings of diameters spanning 1, 3, 7, and 14 degrees of visual angle were presented at the center of gaze in a block design. The paradigm was identical to the one we used previously^63^. We obtained the following data: W, 6 runs, 1 session; Je, 6 runs, 1 session.

Third, to localize regions selective for faces, objects, and colors in the two animals, we ran a standard functional localizer using blocks of natural movie clips ^64^. The blocks included faces, bodies, places, objects, and scrambled objects, which were presented separately in color and in grayscale. The paradigm was identical to the one used previously^64^. We obtained the following data: W, 31 runs, 2 sessions; Je, 29 runs, 4 sessions. Runs with lower than 75% fixation were omitted from analysis.

### fMRI Data analysis

#### fMRI Acquisition and preprocessing

The fMRI experiments were conducted with a Siemens Prisma 3T scanner at the Neuroimaging Facility (NIF) of the National Institutes of Health. We used custom-built eight-channel surface coils kindly made by Azma Mareyam and Larry Wald at Massachusetts General Hospital, and by Charles Zhu and Frank Ye of the NIF. These coils provide full brain coverage. **Fig. S10** shows the signal to noise ratio (SNR) across the cortical surface, computed as the average across the time course for each voxel in the brain divided by the average of the time course of a noise ROI drawn outside of the brain.

Image registration and motion correction were done using the Advanced Normalization Tools (ANTs) register toolkit. All scans were collected with alternating HF (head-foot) and FH (foot-head) encoding directions, except for the active task functional scans, which used FH encoding. Anatomical T1 scans were collected with 0.35mm isotropic voxels. Individual T1 images were intensity normalized, affine co-registered, and averaged to provide a reference anatomical image for each subject, following routine procedures ^63^. Brain masks and white matter masks were automatically generated using FREESURFER (http://surfer.nmr.mgh.harvard.edu), then manually edited. Cortical surfaces were also created using FREESURFER. Functional scans were collected with 1.2mm isotropic voxels and a 17ms echo time. Functional scans were thermal denoised using the NORDIC tool, distortion corrected using FSL top-up (https://fsl.fmrib.ox.ac.uk; HF and FH spin-echo images were collected for each scan session), and corrected for sphinx position. For a few functional localizer and passive-viewing-task scan sessions, spin echo images of opposite encoding direction were not collected. In these cases, top-up correction was omitted and nonlinear registration of the session to standard functional space was done instead. For the active task, slice timing correction to the middle time point of each TR was performed on the raw EPI data.

The median image of all functional scans within a session was used as the session representative volume, and all other images in the session were linearly motion corrected to this target.

Additionally, we chose the median volume of all session representatives for each subject as a functional target, which defined a subject specific functional space. A mapping to this standard functional space was constructed by affine registration of each session representative to the functional target. The session functional data (EPI scans) were then projected to the subject’s standard functional space. The functional target was nonlinear SyN registered to the subject’s anatomical MRI. This procedure allowed us to run GLMs, compute contrasts, and run the decoding models in each subject’s native functional space. This is desirable because multivoxel decoding relies on high-spatial-frequency information, which is degraded by smoothing or interpolation^88,89^. Results (z-contrasts, accuracy maps, etc.) were projected to the anatomical reference, and then to the cortical surface. To test replicability, we analyzed each subject’s data separately for all experiments.

### Region of interest (ROI) definition and brain segmentation

As described below, our primary analyses were performed over the whole brain and did not require pre-defining regions of interest. Nonetheless, to relate our findings to other work and to standard atlases, we generated regions of interest defined by anatomical and functional parcels. These ROIs included retinotopic areas (V1, V2, V3, V3a, V4, MT), partitions within inferior temporal cortex (posterior, central, anterior: pIT, cIT, aIT), the temporal pole (TP), partitions of prefrontal cortex (orbital, ventral, dorsal: oFC, vFC, dFC), intraparietal sulcus (IPS), dorsal superior temporal sulcus (dSTS), the medial temporal lobe (MTL), cingulate cortex (CC), operculum (oprc), primary auditory cortex (A1), the hippocampus (hpc), insula (insl), striatum (str) and superior colliculus (SC). These parcels were defined for each subject using a combination of anatomical landmarks, the D99 macaque atlas ^90^, and functional localizers.

### Decoding analysis of the active-task data

We used fMRI data from the active task to train separate decoders to classify shape and color identity, and then to evaluate these decoders across features. We ask: can a classifier trained using neural data elicited by concept colors, decode the color associations of the concepts using neural responses elicited by concept shapes? And vice versa, can a shape classifier trained using data elicited by concept shapes, decode the shape associations of the concepts using data elicited by the concept colors?

Recall: a single trial in the active task consisted of 7 TRs and each TR was 3 seconds (**Fig. 2a**). The first 3 TRs presented a colored blob or colorless shape (the “cue”) during which the animal was required to maintain fixation. The fourth TR presented four choice options and the animal selected one using an eye movement to receive a drop of juice. The last 3 TRs were blank gray screens, providing a 9-second inter-trial interval before the presentation of the cue for the next trial. The classifiers were trained using data from the cue TRs, not the choice TR since the choice TR contains the feature (color or shape) associated with the cue (shape or color) that we aim to decode.

Conventional fMRI analysis involves constructing a General Linear Model (GLM) with a design matrix that accommodates all known factors that may explain measured brain signals. The design matrix typically incorporates a hemodynamic response function (HRF) that temporally deconvolves the fMRI signals—this deconvolution is useful because the fMRI signal is delayed relative to the sensory stimulus. But adopting this approach in the present context raises an obvious concern: activity attributed to a cue TR might partially be caused by the subsequent choice TR. To completely circumvent this possibility, we did a conservative analysis: we constructed the design matrix without an HRF for the cue TRs. In other words, each cue TR-regressor is an impulse function active for that single TR only (**Fig. 2a**). The cost of this decision is that any signal elicited by the cue TR that is not coincident with a cue TR will not be captured—because of the hemodynamic delay of the MION signal, there is some signal elicited by the cue that would emerge after the cue TR. We mitigated this cost by including three consecutive cue TRs in the task. The MION signal peaks between 2.5 and 3 seconds after stimulus onset ^52^, so we expect strong cue-related signal in the second and third cue TRs of each trial which would be mostly attributed to brain activity in the first and second TRs. We do not expect strong cue signal during the first cue TR, so although this cue TR is included in the design matrix, we did not train the classifiers using signals assigned to it.

Conventional fMRI analysis also typically involves building a design matrix that includes confound regressors that allow the Generalized Linear Model (GLM) to regress out scanner drift, body and head motion, eye movements, and other non-specific high variance confounds. We accommodate these regressors into our design matrix too. Confound regressors included cosine drift regressors with a high frequency cutoff of 0.03 Hz, two CompCor regressors^91^, median fixation distance over the TR, and six motion regressors accounting for translation and rotation, all convolved with the standard MION HRF^52^.

Finally, we constructed our design matrix to incorporate regressors for the residual cue signal that emerges after the end of the third cue TR (estimated with the standard MION HRF) and the choice TR (also estimated with the standard HRF) (**Fig. 2a**). The regressor for the choice TR ensures against the unlikely possibility that the neural signal measured during the cue TRs are confounded by signals elicited by the choice TR of the prior trial. This possibility is unlikely for two reasons: first, because there was a 9 second inter-trial interval, during which the MION signal would substantially decay; and second, because the sequence of trials is counterbalanced and random, so a preceding choice TR is uncorrelated with a subsequent cue TR. We confirmed that this is, indeed, the case: the portion of trials where the previous correct answer is the same as next cue is at chance (monkey W; median 0.003, 95% CI [-0.004, 0.010], monkey Je 0.000, 95% CI [-0.006, 0.005]). Chance was determined by the portion of pairs of trials that fill each case over the total number of possible pairs of sequential trials.

We recognize that our design matrix is somewhat unusual, but it enables us to be sure that the classifiers depend solely on neural signals attributed to the cue TRs and still controls for known confounds such as scanner drift and body motion. The procedure for constructing our design matrix is as follows. On each run, we use a separate HRF-regressor for the cue and choice TRs of each trial. As previously mentioned, we also define a TR-regressor (an impulse function that has value 1 for a single TR and 0 everywhere else) for each cue TR. We then zero the cue HRF-regressors at every TR where this is an active TR-regressor. This has the effect of modelling the TRs where the cue is present as a single timepoint (the TR-regressors) and preserves the cue’s impact on future non-cue TRs as the remaining non-zeroed tail of the cue HRF-regressors. The separate choice HRF-regressor for each trial captures the unique signal from the four presented choices as well as any signal from the reward (**Fig 2a**).

After fitting the GLM, we took the volume matrix of coefficients for the second and third cue TR-regressors as the input data to the decoding model for that trial (the target was the identity of the cue). Once trained, the classifiers were used to decode feature identity: they tested using data elicited by the same feature (color or shape) as used to elicit the data for classifier training (but where the training and testing data were independent). This analysis answers the question: from a pattern of neural activity elicited by a given cue (a color or a shape), can the classifiers determine the cue identity (which color or which shape)? We refer to these tests as “train on shape; test on shape” or “train on color; test on color”. The classifiers were also used to decode across features: they were trained on data elicited by one feature (color or shape) and tested using data elicited by the other feature (shape or color). The analysis answers the question: from a pattern of neural activity elicited by a given cue, can the classifiers determine the identity of the unshown feature linked by an object concept to the identity of the cue? We refer to these tests as “train on shape; test on color” or “train on color; test on shape”. The procedure of cross-validation is described below in the section *Validation of searchlight models*.

### Layered Searchlight Decoding Model (LSDM)

For the main analysis, we developed a new method we call the Layered Searchlight Decoding Model (LSDM, **Fig. 2b, 2c**), which is inspired by convolutional neural networks. We applied the LSDM to data in-volume (i.e. in 3-dimensions) to determine in a data-driven manner where and how color, shape, and putative color-shape conceptual representations are distributed across the whole brain. The LSDM is like a convolutional neural network in that it is composed of small stacked linear filters with an increasing input domain at each layer. But unlike a convolutional network (e.g. for image classification), where the same filters are applied across all input data, the LSDM learns a filters at each spatial location. Having the flexibility to learn different filters across spatial locations allows the model to discover the extent of decoding at each local region of the brain.

A modified layer 𝐻 was defined that applies overlapping cubic kernels of size 𝑘 with different learnable weights at each location. This is a convenient way of packaging many separate linear classifiers acting at different locations. Like a standard convolutional layer, each 𝐻 has an input and output channel dimension, 𝑐_!"_ and 𝑐_#$%_. By putting L of these layers in sequence, the LSDM obtains an output at each index of the input with access to only a cube of size 𝐿𝑘 − 𝐿 + 1; 𝑐_#$%_ is equal to the number of classes for the output layer; and 𝑐_!"_ is equal to the number of TRs per trial for the input layer. The intermediate channel dimensions, 𝑐_&_, are hyperparameters that scale the number of model parameters. These were set to 3. (This number accommodates the three dimensions of color space.) We set 𝐾 to 2 and 𝐿 to 4, so the output of each searchlight patch is fully determined by the values of a 5x5x5 cube of voxels (6x6x6 mm). If 𝑘 were set to 5 and 𝐿 were set to 1, we would obtain a standard searchlight over a 5x5x5 cube of voxels.

The LSDM provides three advantages. First, its parameters are shared between adjacent filters, resulting in significantly smaller models compared to a conventional searchlight. This allows a consumer grade GPU to fit the full model over the whole volume with reasonable batch sizes and is much faster than existing searchlight implementations. Second, the LSDM has fewer parameters compared to a conventional searchlight and low dimensional bottlenecks between layers, which improves performance when applied to noisy data typical of fMRI. Third, operations such as batch normalization, dropout (randomly zeroing activations), and nonlinear activation functions (e.g. ReLU) can be applied between layers of the LSDM, allowing modelling of more complex relationships than is afforded with a conventional searchlight. In the current report, we apply batch normalization at each spatial location between layers. No nonlinear activation functions where used, but it is conceivable that they could improve performance of the LSDM even further relative to a standard searchlight. The output of the standard (non-layered) searchlight is compared to the LSDM using simulated data and real data in **Fig. 3b-j**. The LSDM achieves better performance because it functions as a method for regularization. We provide an analysis of the LSDM using simulated data sets because these data sets are computationally less demanding to analyze than real fMRI data, which facilitates systematic assessment of the approach given different kinds of noise. It will be fruitful to conduct in future work a fuller assessment of the conditions for which the LSDM approach is especially advantageous.

We were motivated to develop the LSDM to solve two problems. First, we needed a method that could rapidly train and test models. Training the LSDM requires about 1/5^th^ of the memory of an equivalent logistic searchlight and thus is more easily parallelized, i.e. much faster. Second, and more critical, we wanted to represent the stimuli in a shared low-dimensional latent space, which is not afforded by a standard logistic or SVM model that fits a separate weight vector for every class. The representation of input data in the final hidden layer of the LSDM (the LSDM latent space) is attractive because it has linearly separable representations of each class on a low dimensional manifold (See *Computing ROI dissimilarity matrices from LSDM states* section for more details). The LSDM latent space has three dimensions, corresponding to the number of hidden channels (𝑐_&_ = 3) for each searchlight patch.

Having a shared low-dimensional space is useful to test for structure of the representational space (see section below, *Comparing task representations with color space geometry*). In the LSDM, all data are projected to a low-dimensional manifold constructed such that the shape or color features of the concepts are optimally decodable. Like other neural population analyses where neurophysiological data or fMRI data are projected to task-relevant axes before analysis of representations or dynamics, the LSDM is useful because it is not impacted by off-manifold task-irrelevant signals that affect the distances between classes, which can obscure recovery of the underlying geometry.

An additional advantage of the LSDM, and how we define its object function, is that it readily accommodates the fact that the trials were divided up into different sets of runs, A & B (see active choice task methods). There will be signal and noise along dimensions that are not consistent between individual shapes or colors simply because the data were collected in separate sets of runs—such differences could include different noise structures or set context, which would be unrelated to the individual trials, and critically orthogonal to LSDM latent space because they are not useful for decoding between items within set and luminance level. The LSDM latent space therefore minimizes the influence of possible nuisance differences in the sets.

All code for the LSDM (and other analyses) is available via github.com/NEI-LSR/ColorShapeConcepts.

### Validation of searchlight models

Metrics computed via the searchlight model were validated using 8-fold cross validation (CV). Reported error bars are 95% confidence intervals (CIs) computed by bootstrapping 1000 times over each CV measure (see **Figs. 2, 4, S4, S6**), following common practice. We note that, for the identity-decoding conditions, this estimate of variability is approximate, because separate CV measures are not fully independent: the train sets of the C.V. folds will overlap, even though the test set and train set for each fold are entirely independent^92,93^. We determined that procedures such as nested cross validation^94^, which could produce unbiased confidence intervals on the identity decoding, were unnecessary given the large effect sizes we observe, the added computational cost, and their need for normality assumptions. For all cross decoding and error-decoding conditions, the test data folds are truly independent of the data used to generate the models, since they are never included in any fold of the model’s training data, so the error bars are true estimates of the variability in the underlying signals. Each fold of the test data set was evaluated by a model trained on ^’^ of the total data, and the trained models for the cross decoding and identity-decoding analyses were the same. We could have performed the cross-decoding analysis with a model trained on ^(^ of the data, since the training and testing data are entirely independent—this would offer slightly better estimates of cross decoding but would compromise the direct comparison of the identity and cross decoding results (see **Figs. 2**, **S3, S4**).

For cortical surface maps showing the decoding results (**Figs. 2, S3**), the FDR-corrected chance threshold was computed via permutation testing for the data for each individual monkey on each animal’s own cortical surface. For both color and shape input data, each searchlight patch for each trained model (one model for each of 8 CV folds) predicted a different random permutation of the class labels over its corresponding test set (1/8 of all data). This was done 1000 times. The accuracy results for each iteration were then projected to the cortical surface of their respective monkeys, since this projection involves averaging accuracies over multiple searchlight patches. All results (values at each surface vertex for each iteration over both monkeys) were concatenated and we found the 95^th^ quantile of the resulting distribution (**Fig. S3**, color-bar).

This value is chance threshold, FDR-corrected, for all surface maps, which enables a direct comparison of the decoding results across monkeys and classifiers. The standard notion of chance for a balanced dataset is the mean of the distribution and sits at 1/3^rd^ of the color scale bar, as expected. We show the distribution of decoding accuracies that were below threshold for completeness.

### Region of Interest Analysis

Given a trained searchlight model and a region of interest (ROI, defined to be a contiguous set of voxels), 𝑅, predefined without any knowledge of decoding, the naïve way to get accuracy over 𝑅 is to take the average searchlight accuracy for the individual searchlight spots contained by R. But this would be misleading because it would include uninformative predictions from searchlights within the ROI that provide no signal, reducing measured accuracy and compromising the ability to compare performance across ROIs (since different ROIs could contain wildly different numbers of uninformative searchlights). So instead, we learned the stacking weights 𝑊^-^ over the whole brain to take a weighted average of the predicted class distribution 𝑐_!_ ∈ 𝐶 from each of 𝑛 searchlight spots. 𝑊 can be thought of as a probability distribution over 𝐶, and 𝑐^∗^ = ∑^"^ 𝑤_!_ 𝑐_!_ where 𝑐^∗^ is a probability distribution over the classes.

After training, 𝑊^-^ is approximately the maximum likelihood solution to the objective given 𝐶. Any subset 𝑢 ⊆ 𝐶 has a matching subset 𝑣 ⊆ 𝑊^-^, where 𝑤_!_ ∈ 𝑣 ⇔ 𝑐_!_ ∈ 𝑢. By renormalizing 𝑣, we obtain the maximum likelihood estimate solution for the objective over 𝑢, and a weighted sum yields the probability of each class for that subset of searchlight spots. We tested the combined model (trained searchlight and weights) on held out data, giving a single accuracy metric in each ROI. These weights were also used in the representational geometry analysis described in the section *Comparing task representations with color space geometry*. Note that this method is agnostic to the particular searchlight model used, so long as it produces some distribution over classes at each spot.

### Decoding objective function

The output of the searchlight is a predicted score for each class. Normally, a multinomial logistic classifier would compute a distribution over the classes (e.g. the softmax of the scores) and the objective function would be the cross entropy of this predicted distribution and the true class distribution. In the present work, not all class comparisons are valid, for two reasons. First, the colors were presented at two luminance levels. In order to mitigate the effect of luminance we only allow within-luminance-level class comparisons. Second, in order to effectively counterbalance the stimuli in each MRI run we had divided the overall set of shape / color pairs into two groups (group A and B), each with three light and three dark colors (**Fig. S2a,** see active choice task methods). We only compare exemplars that are in the same set. These considerations result in four groups of conditions (group A light, group B light, group A dark, group B dark) each with three shape / color pairs. The cross entropy and accuracy are calculated only within group, so chance is 1/3. This strategy only considers meaningful class comparisons, but the LSDM still jointly represents all 12 class pairs in the same space until the last readout layer and thus can exploit shared structure. Besides improving decoding accuracy, this shared latent space is especially important for analyzing the representational geometry of shapes and colors (**Fig. 4a**).

As described above, the animals performed slightly unbalanced numbers of the different trial types (especially for subject Je, where fewer runs of group B were collected than group A). The dataset was balanced by oversampling (after train / test splits) but classes with fewer original examples would still be more prone to overfitting. This problem, as well as any other difference in classification difficulty across classes, was mitigated by dynamically adjusting the weight given to each class based on its difficulty, as done by others^85^.

### Simulation to compare LSDM and standard searchlight

We built a simulation to compare the performance of the LSDM and standard searchlight (**Fig. 3**). For a sketch of the difference between the LSDM and standard architecture see **Fig 3a**; the LSDM searchlight has fewer parameters (**Fig. 3g**) because some parameters can be shared between adjacent filters. In the simulation, there were two modalities, corresponding to the two features of the object concepts, S and C, each of which had 12 classes. Classes with the same index in S and C were “associated”. Each data example was a 16x16x16 volume. We defined two 5x5x5 regions of this volume that could carry signal about class identity, one which was in the upper left (UL region) and one in the lower right (LR region) of the first two dimensions of the volume, both centered on the middle (index 8) of the third dimension (**Fig. 3b**). The UL-only region carried information about S, whereas the LR region carried some shared information about both S and C. As in the actual experiment, we divided the classes into two luminance levels (corresponding to light and dark colors) and two groups (alternating colors from each luminance level) (**Fig. 3e**).

The class identity signal was the sum of a circular and high-dimensional signal. The circular signal was defined by placing the 6 classes in the same luminance level evenly around the unit circle and then projecting them to a plane in a 125-dimensional space via a randomly chosen orthonormal transform (since the UL and LR regions were 5x5x5). Examples belonging to the “dark” luminance level were then offset by 1 along a random axis orthogonal to the plane of the circle. In the LR region, the circular signal was shared between associated classes from S and C. The high-dimensional signal was designed to orthogonalize the 12 classes and thus is simply an orthonormal transform of their one-hot encoding into a random 12-dimensional subspace of each 125-dimensional (5x5x5) region. The high dimensional signal was not shared between S and C in either region and was scaled by a factor of 1.5 (so that cross and identity decoding accuracy were similar to the real data). Finally, random 4096 (16x16x16) dimensional isotropic Gaussian noise with spread of 1.5 was added over the whole volume for every example generated independently (regardless of class label). In the structured noise case only, independently drawn 4096-dimensional signal (from an isotropic gaussian with spread of 0.3) was added to Group A and Group B data, in this case noise structure is dependent on class identity. The hyperparameters were selected to give unsaturated performance on dataset sizes similar to our real data. We empathize that this simulation is not intended to provide an exhaustive comparison of different searchlight methods, but to explore why the LSDM is well suited for our case. It’s likely that other searchlight methods could perform better under different conditions. For instance, if the true signal is very high dimensional or changes rapidly over space, it is expected that a standard searchlight would perform better.

We attempted to decode from 5x5x5 searchlight patches, just as in the actual experiment. We expected to see cross decoding (train S, test C) only in the LR region. We plotted heatmaps of cross decoding accuracy (difference from chance) for searchlight patches centered on the middle of the third dimension of the data volume for both the LSDM and standard logistic searchlight (**Fig. 3c**), as well as heatmaps of the stacking weights (see *Region of Interest Analysis* section) (**Fig. 3c**) for an input dataset size of 180 examples per class. To quantify accuracies, we defined two non-overlapping ROIs (**Fig. 3b**): ROI 1 that contains the UL region and ROI 2 that contains the LR region. For individual identity and cross decoding accuracy metrics, we report the combined accuracy over ROI 2 (see *Region of Interest Analysis* section). We did this for a variety of input dataset sizes (60, 120, 180, 300, 420, and 660 examples of each class for both S and C; **Fig. 3d**). Varying the dataset size impacts the ease of test-set decoding because noise was independent for each example, so smaller dataset sizes would be overfit. All accuracy metrics shown were 8-fold cross validated as described in previous sections for the analysis of real fMRI data. The LSDM consistently performed better at both identity and cross decoding for all dataset sizes. These plots results are for the unstructured noise case, but accuracy in the structured noise case is similar – only within group comparisons are considered, just as for the real data (see Layered Searchlight Decoding Methods). However, the structured noise is deleterious to the recovery of the underlying class geometry using standard RSA techniques. With unstructured noise, the ground truth geometry (**Fig. 3e**) is recovered well either by RSA in the LSDM latent space (blue dotted line) or with standard RSA on the raw data (grey dotted line), though the LSDM does slightly better (**Fig. 3f, top)**. With structured noise RSA in the LSDM latent space still recovers the ground truth geometry, but standard RSA achieves only very weak correlations (**Fig. 3f, bottom**). We believe this is because the LSDM recovers the low dimensional decodable space where class geometry is preserved. This may explain why standard RSA does not recover reliable correlations with color space on our real data (**Fig. 4**). Simulation code is available at https://github.com/NEI-LSR/ColorShapeConcepts.

### Determining the structure of the representational space: Computing ROI dissimilarity matrices from LSDM states

We wanted to be able to compute the representational geometry of color and shape stimuli in the LSDM’s final latent space, because it is 1) low dimensional (𝑐_ℎ = 3 dimensions), and 2) it must have linearly separable representations of the classes, since the final transform is a linear readout. These properties make the LSDM latent space useful for analyzing representation geometry, as described above.

Let each 𝑥_!_ ∈ 𝑋 be a correct trial, that is a unique presentation of a shape or color, and 𝑇 = {𝑡_!_: 𝑡_!_ ∈ ℕ, 0 < 𝑡_!_ ≤ 12}. Let 𝑉 be the set of all voxels, or more generally, all integer locations in the affine functional MRI space 𝑆. Our goal is to compute a distance matrix 𝐷_1_ over ROI 𝑅 ⊆ 𝑉. Moreover, we will extract these representations from an LSDM model trained on (𝑋, 𝑇) with 𝐿 layers 𝐻^2^, 0 < 𝑎 ≤ 𝐿 and kernel size 𝑘. Let 𝑗 ∈ ℕ, 0 < 𝑗 ≤ |𝑅| be an index for each location in 𝑅. 𝑈^2^ denotes the full set of outputs from layer 𝐻^2^. Each 𝑢_3!_ ∈ 𝑈^2^ is a ℝ^4^ vector at each location 𝑗 for each trial 𝑖. Let 𝑇_5_ = {𝑖: 𝑡_!_ = 𝑏, 𝑡_!_ ∈ 𝑇, 0 < 𝑏 < 12}. We obtain an embedding for each class in each location by letting

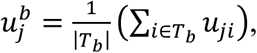

which gives an embedding of each class 𝑏 at each location in space 𝑗. Taking, 𝑓(𝑥, 𝑦) to be the Pearson correlation distance between two vectors, we construct a distance matrix 𝐷_3_ at each location.

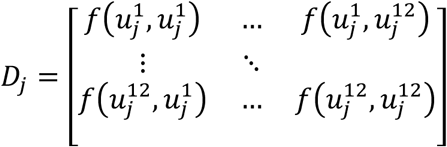

We now have a single distance matrix 𝐷_3_ for every location in 𝑅. We want to combine these in an intelligent way to obtain 𝐷_1_, a single distance matrix for 𝑅. We already have weights 𝑊, from training, which give the optimal way to combined predictions from layer 𝐻:. Let 𝑙_3_ ∈ 𝐻: be an individual filter at location 𝑗. The input to each 𝑙_3_ in then a size 𝐾^;^ subset of the output of the previous layer 𝑈_:<)_, let this set of locations be called 𝐼_3_. Let 𝐷^b^_=_ be the average distance matrix over 𝐼_3_,

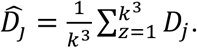

Note that this is equivalent to running a (𝑘 × 𝑘 × 𝑘) uniform filter over the distance matrices computed on the outputs of layer 𝑚 − 1. We think of each 𝐷^b^_=_ as the average representation that is sent to the final layer that will give a linear readout of class predictions. We construct 𝐷_1_ as the average of all 𝐷_3_ weighted by 𝑊, normalized over R,

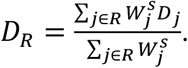

### Determining the structure of the representational space: Comparing task representations with color space geometry

The colors of the object concepts evenly sampled the hue dimension of CIELUV color space, at two luminance-contrast levels ("light" and "dark"). If a representation of color or associated shape in the brain matches the geometry of color space, items within a luminance level should be arranged in a circle with the same ordering. We will call the canonical ranking of six evenly spaced items around a circle 𝑂^4^.

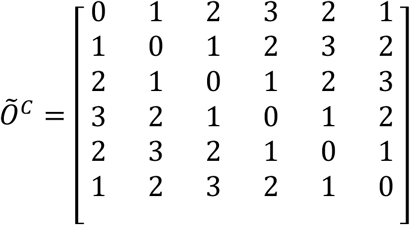

𝑂^D^ is the upper triangle of 𝑂^fD^, not including the diagonal. We let 𝐷^E^ and 𝐷^C^ be size (6 × 6) sub-matrices of 𝐷_1_ corresponding to light and dark elements respectively. The corresponding 𝑂^E^ and 𝑂^C^ are the rank of elements in the upper triangles of 𝐷^E^ and 𝐷^C^. Then we define the correlation metric,

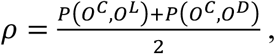

where 𝑃(𝑥, 𝑦) indicates the pearson correlation of two vectors. Note that the pearson correlation of two sequences of ranks is equivalent to the spearman correlation of the vectors that were ranked in the absence of ties. This metric 𝜌 is a measure of how similar a relevant representational geometry is in an ROI 𝑅 to color space. We show plots of correlation for light and dark color-shape pairs separately (𝑃(𝑂^D^, 𝑂^E^) and P(𝑂^D^, 𝑂^C^) respectively) in **Fig. S6b**.

We took an LSDM trained on color stimulus data to compute 𝜌 on both held out shape and color test sets. Using the same k-fold cross validation procedure used for accessing decoding performance, we obtained k measures of 𝜌 for each ROI for both color and shape test sets. We plot both mean 𝜌 and 95% CIs computed by bootstrapping 1000 times over 𝜌. CIs that are above 0 indicate that a representation is positively correlated with color space.

What should we take to be a significant 𝜌? Since each correlation is dependent on the ordering of only 6 items, we must account for the fact that a random ordering might correlate with color space by chance. By generating 10000 random orderings of six items and then computing 𝜌 with respect to color space, we determined the correlation threshold that is not exceeded by chance 95% of the time. This is the chance level (dotted line) in plots of 𝜌 (**Figs. 4a, S6).** When combining multiple measures, 𝜌 was averaged on each iteration, which explains why the threshold is slightly higher in **Fig. S6b** (two independent measures of 𝜌) than in **Fig. 4** (four independent measures of 𝜌).

### Determining the structure of the representational space: without the LSDM

The above calculation of 𝜌 requires a trained LSDM model and stacking weights 𝑊^-^. In classic RSA, dissimilarity matrices within an ROI are computed over condition vectors in voxel space^95^. We performed this standard analysis on each ROI (**Fig. S6c**), using Pearson dissimilarity as the metric. In order to take advantage of the learned weights 𝑊, (i.e. how much each searchlight patch should matter), we constructed a dissimilarity matrix 𝐷_3_ for every (𝑘 × 𝑘 × 𝑘) subset of voxels, and then we took the weighted sum of 𝐷_3_ over R. So,

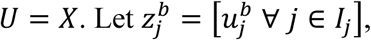

i.e. the concatenation of all 𝑢^5^ over a (𝑘 × 𝑘 × 𝑘) cube of data. Then,

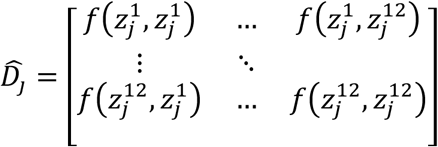

𝐷_1_ can now be computed and compared with color-space geometry as above. This procedure was intended to be as similar as possible to the one in the main text, except distances matrices were computed on the raw data for each patch instead of in a trained model’s latent space.

Results from this alternative procedure are compared to results using representations extracted from a trained LSDM, see **Fig. S6**.

### Computing incorrect trial decoding

We defined a valid trial on the active fMRI task by two criteria: during the stimulus block, the monkey had to have been fixating within 1 degree visual angle (dva) of the fixation dot, and during the choice block, the monkey must have fixated one of the four options for at least 1000ms (the “choice”). A correct trial was one where this choice was correct given the stimulus; all other valid trials were deemed incorrect.

On the incorrect shape and color trials, we used the pretrained shape and color identity models to predict the true stimulus identity. **Fig. 4b** shows pooled shape and color stimulus decoding on incorrect trials. We also tested whether we could predict incorrect choices. To do this we used cross decoding (e.g. the model trained on correct shape identity trials predicts incorrect choices on color trials) since we are interested in whether shared representations are related to motor output. **Fig. 4c** shows pooled shape to color and color to shape cross decoding of the monkey’s choice on incorrect trials.

### Passive viewing task 1: colorless color-associated shapes versus colorless non-color-associated shapes

Animals were presented with blocks of stimuli (**Fig. S7a**) while collecting fMRI data. Each run was separately fit with a GLM, with the stimulus conditions as regressors of interest: colored shapes, colorless color-associated shapes, colorless non-color-associated shapes, and shapeless colored blobs. Confound regressors included cosine drift regressor (high frequency cutoff defined as the reciprocal of twice the longest time period between two blocks of the same condition, 0.003 Hz), two CompCor regressors^91^, median fixation distance over the TR, and six motion regressors accounting for translation and rotation. All regressors were convolved with the standard MION HRF^52^. A lag-1 autoregressive (AR1) noise model was used when fitting the GLM.

To compare fMRI responses to color-associated shapes and non-color-associated shapes, we computed standard z-score contrast maps. All z-score contrast maps were computed in-volume as the mean contrast difference divided by the contrast standard error, using the beta weight differences and their corresponding variances from the GLM. The projections of these maps onto the cortical surface are shown in **Fig. 5a**, right; all z-score color bars are symmetric about the gray point. We additionally computed a selectivity index for this contrast. This metric normalizes the difference between two conditions by the additive response to both conditions for each voxel on each run, helping to account for different overall responsiveness and signal-to-noise ratios across the brain. Selectivity was calculated as

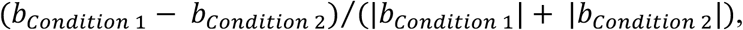

where 𝑏 refers to the beta coefficient for that condition from the GLM fit to a given run. So, the color-associated shape selectivity would be:

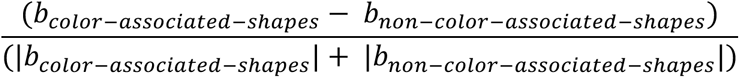

This gives a selectivity value for each voxel in the brain on each run. The boxplot in **Fig. 5a** quantifies selectivity in ROIs using the parcels described in *Region of interest (ROI) definition and brain segmentation* with two additional constraints. First, analysis of responses of retinotopic areas (V1, V2, V3, and V4) was limited to voxels representing the fovea and parafovea using a mask generated by a GLM analysis of the eccentricity localizer experiment. The mask was created by contrasting responses to the two more central stimuli with the responses to the two more peripheral stimuli (one-sided, p<.01, Bonferroni corrected for number of conditions). We did this because the visual stimuli in the passive tasks were presented at the center of gaze within the central 3 degrees of visual angle. Second, for all ROIs analyzed, we included only voxels that were visually responsive, defined as having a different response from baseline to any of the stimulus conditions of interest in the paradigm, i.e., color-associated shapes, non-color-associated shapes, colored blobs, or colored shapes; one-sided, p<.01, Bonferroni corrected for number of conditions. To get one value per ROI, the selectivity was averaged per run over voxels in the ROI, then bootstrapped across runs. The error bars in **Fig. 5a** show 95% confidence intervals computed from the bootstrapped distribution.

### Passive viewing task 2: color-shape combinations congruent versus incongruent with learned object concepts

Animals were presented with blocks of stimuli (**Fig. S7b**) while collecting fMRI data. Other analysis details were the same as for passive-viewing task 1, described above. The only differences were as follows. For the GLM, (1) the regressors of interest were the congruent and incongruent blocks, and (2) the cosine drift regressor used a high frequency cutoff of 0.005 Hz. For the quantification, (1) the contrast was between fMRI responses to incongruent object and congruent object blocks, with incongruency selectivity defined as

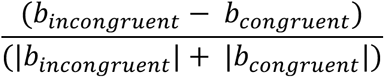

and (2) visually responsive voxels were defined as those having a different response from baseline to incongruent objects or congruent objects; one-sided, p<.01, Bonferroni corrected for number of conditions.

Comparing topography of the areas recovered in the two passive-viewing experiments (areas implicated in color perception and areas implicated in perceptual color-shape binding).

We examined the spatial relationship between signals about color perception and signals about color-shape binding by comparing the magnitude of color-associated shape bias and incongruency bias along the posterior to anterior (PA) axis of the brain. We did so on the cortical inflated surface rather than in-volume, taking advantage of the clear posterior to anterior progression generated by unfolding the cortex (in volume, there is no clear path to follow from V1 to TP). Each inflated surface is defined by vertices, which each contain information from multiple voxels in the volume; e.g., the z-score contrast between color-associated shapes and non-color-associated shapes at a given vertex is a weighted average of the contrast values at the voxels that project to that vertex. For each inflated hemisphere, we isolated the portion of the hemisphere that encompassed V1 through the temporal pole. The distances from V1 to TP differ across the four hemispheres, so to compare and combine across the four hemispheres, we binned vertices into 65 equally spaced bins along the PA axis of each hemisphere separately. To ensure this equal spacing, we spaced bins using geodesic distances, that is, distances calculated along the surface rather than a straight-line axis outside the brain. This normalized distance is represented on the x-axis in **Fig. 5c**, with each data point corresponding to one bin. The y-axis plots the same z-score metric as the cortical surface maps in **Fig. 5a** and **5b** but computes it for each bin of vertices rather than each voxel in the volume. So, within a bin and for each hemisphere separately, the contrast difference and contrast variance were each averaged per run over all vertices in the bin, for both contrasts we were interested in (color-associated minus non-associated shapes and incongruent minus congruent objects). We then concatenated all examples (paired contrast difference and contrast variance) across hemispheres and runs that corresponded to the same bin, giving each bin 74x2=148 examples (number of runs x number of hemispheres; each subject’s two hemispheres were treated separately). We bootstrapped across examples, computing a z-score for each bin on each resample by dividing the mean contrast difference by the contrast standard error. In **Fig. 5c** solid lines plot the mean z-score, and the shaded regions cover the 95% confidence interval. In the interest of mapping out the broad pattern along the PA axis, we did not exclude peripheral voxels from this analysis.

### Testing for color responses elicited by colorless color-associated shapes within color-biased regions

We computed the percent signal change difference between responses to color-associated and non-associated shapes (“color-associated shape bias”) in color-biased and non-color-biased voxels comprising V4, pIT, cIT, and aIT. Percent signal change was calculated by dividing the beta coefficients for color-associated shapes and non-color-associated shapes each by the mean baseline response across the whole brain and multiplying by 100, for each run and voxel separately. The difference between conditions was taken and then averaged per run over voxels in each color-biased and non-color-biased region, giving one color-associated shape bias value per run per region.

The color-biased and non-color-biased regions of V4 and IT were defined using the third functional localizer described above, as described previously^64^. Within each ROI, we considered only visually responsive voxels (responsive to any localizer condition above baseline, one-sided, p<.01, Bonferroni corrected for number of conditions). These were then split into color-biased voxels (those responding significantly greater to all color conditions than all grayscale conditions, p<.01 was used, one-sided, FDR corrected) and all other voxels, yielding the numbers of voxels in each ROI as follows: Je V4: ncolor = 287, nother = 303; W V4: ncolor = 698, nother = 142; Je pIT: ncolor = 202, nother = 447; W pIT: ncolor = 596, nother = 141; Je cIT: ncolor = 245, nother = 374; W cIT: ncolor = 548, nother = 244; Je aIT: ncolor = 31, nother = 316; W aIT: ncolor = 371, nother = 344;

We then subtracted, for each run separately, the color-associated shape bias in the non-color-biased voxels from the color-associated shape bias in color-biased voxels. We did this procedure multiple times, varying the proportion of voxels included in the population of color-biased voxels, from all the voxels (100%), to the top 10% most color-biased voxels. The extent of color-bias was determined by ranking the voxels based on their z-scored contrast difference, comparing responses to color images with responses to grayscale images in the localizer experiment. For this analysis, the number of voxels defined as non-color-biased was fixed across iterations to be the set of visually responsive voxels that were not identified as color-biased in the functional localizer.

### Bootstrapping confidence intervals

In all cases where bootstrapping is used to construct confidence intervals, we used the following procedure. Each of 𝑚 subjects’ data 𝐷_!_ , 0 < i ≤ m was concatenated, then sampled with replacement with the probability of drawing each example,

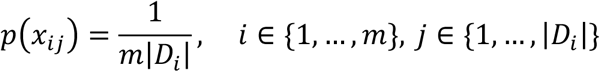

This ensures that on average an equal number of examples from each subject are included in each iteration. The sample mean was taken for each iteration. This was repeated for 1000 iterations, and the distribution of the sample means was used to construct 95% intervals around the final mean over all samples. When bootstrapping is done over raw fMRI data, each example is taken to be a single run.

### Predictive coding model

The predictive coding model is a simple two-layer bi-directional network with symmetric weights. The first “perceptual” layer 𝑃 has a separate feature for each shape and each color. In the example (**Fig. 7a**) there are two possible colors and three shapes, so the perceptual layer is 5 dimensional. The “conceptual” layer 𝐶 has a feature for each concept. In the example (**Fig. 7a**) there are three concept features. The weight matrix 𝑊 is then a 5 × 3 matrix that maps perceptual features onto a concept. For instance, since the first concept feature in the example is a pink hourglass, the first row of the weight matrix will have a 1 in indexes corresponding to “pink” and “hourglass” and a 0 elsewhere. To simulate the accumulation of information over time, activation in both the conceptual and perceptual layer is modeled as a discrete-time iterative update process, where each layer gradually adjusts its activation in response to prediction errors. The rate of this adjustment is controlled by α, a parameter that determines how quickly the system updates its activations. Given input 𝐼 and assuming the identity mapping between the input and perceptual layer the simulation proceeds as follows at time step 𝑡:

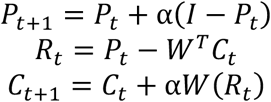

Where 𝑅_%_ is the output from the perceptual layer and input to the conceptual layer at time 𝑡. 𝑅_%_ is plotted over time for different inputs in **Fig. 7b**.

## Data Availability

All data required to reproduce the analyses will be available at https://figshare.com/projects/Datasets_for_The_representation_of_object_concepts_across_the_brain_/255242.

## Code availability

Code for all analyses, including the LSDM and representational geometry, is available at https://github.com/NEI-LSR/ColorShapeConcepts. See the GitHub repository README for directions on how to access and run each analysis. Code for psychtoolbox paradigms defining fMRI stimuli is available at https://github.com/NEI-LSR/ColorShapeConcepts-fMRI-Paradigms.

## Acknowledgements

This research was supported in part by the Intramural Research Program of the National Institutes of Health (NIH, 1ZIAEY000558 to BRC). The contributions of the NIH authors are considered Works of the United States Government. The findings and conclusions presented in this paper are those of the authors and do not necessarily reflect the views of the NIH or the U.S. Department of Health and Human Services. The research was also supported in part by grants from the National Science Foundation (0918064 to BRC) and NIH (R01 EY023322 to BRC). We thank David Leopold and the Neuroimaging Facility at the NIH. Noah Lasky-Nielson provided help running some fMRI experiments. For help training the animals, we thank Shridhar Singh, Kaitlin Bohon, Walid Bendris, and Isabelle Rosenthal. We are grateful to Alex Martin, Chris Baker, Eli Merriam, Doris Tsao, Thomas Nasaelaris, Hendrikje Nienborg, and Marlene Cohen for feedback, and to members of the Laboratory of Brain and Cognition (NIMH) and the Laboratory of Sensorimotor Research (NEI) for discussions. We thank Azma Mareyam and Larry Wald for providing the fMRI coils, and Denise Parker, Hayden Warnock and the outstanding team of animal care technicians and veterinarians at the NIH for expert animal care. We also thank Jeeves and Wooster for years of dedicated service to this project.

## Author contributions

Design of the active-task fMRI experiment (SRL, SD, BRC), fMRI data collection (SRL, HEF, SD, KK), analysis of fMRI data (HEF, SRL, KK), development of the layered searchlight decoding analysis and representational geometry analysis (SRL), generation of figures (SRL, HEF, BRC), writing the manuscript (BRC, SRL, HEF), project conceptualization, oversight, and funding (BRC).

## Notes

https://figshare.com/projects/Datasets_for_The_representation_of_object_concepts_across_the_brain_/255242

https://github.com/NEI-LSR/ColorShapeConcepts

### Competing Interest Statement

The authors have declared no competing interest.

