## Supplementary Figures for "The representation of object concepts across the brain"

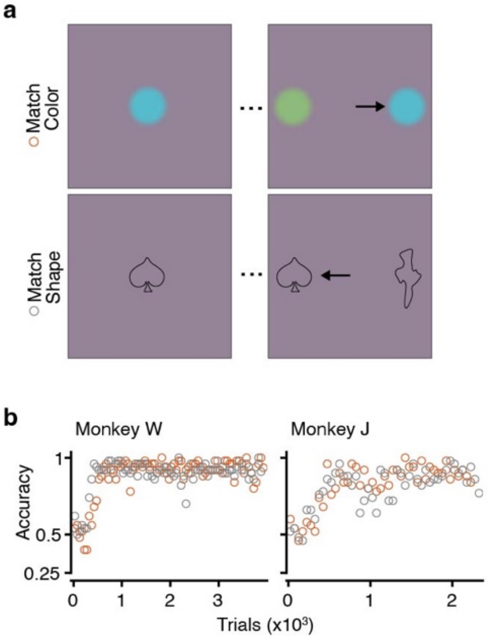


**Fig. S1: Colors and shapes of the object concepts are equally discriminable. a,** Color-to-color and shape-to-shape matching trials included in 2-alternative-forced-choice short-term memory trials. **b,** Accuracy of the two trial types over time; data points are averages of 50 trials. Performance on the two trial types is not significantly different (% correct color-to-color, % correct shape-to-shape, [95% CI], for the last 1000 trials: W, 93.3% [91.7, 94.9], 92.4% [90.6, 94.0]; J, 91.2% [89.6, 92.8], 88.8% [86.9, 90.8]).

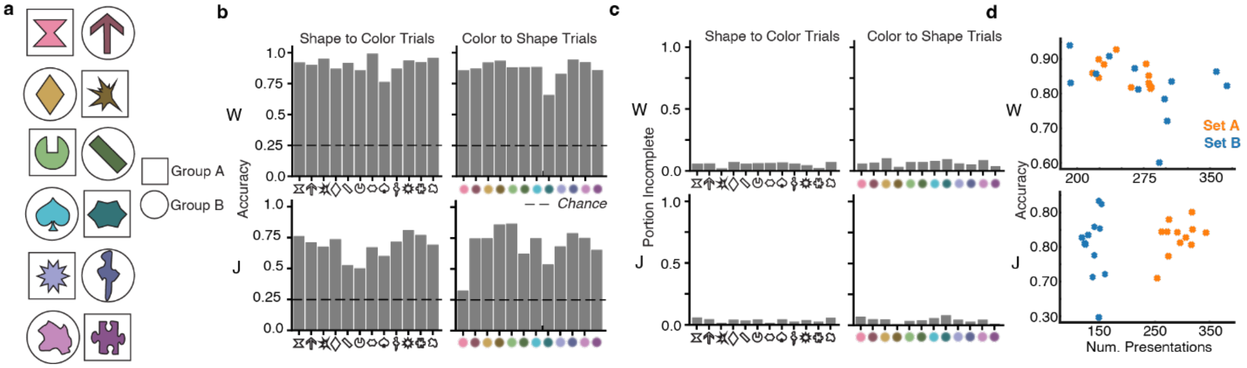


**Fig. S2 Monkey performance during fMRI scans a,** Object concepts tested in the trials in the main fMRI experiment and the group of runs to which they were assigned (see Methods). **b,** Behavioral accuracy for trials performed in the fMRI experiments, separated based on trial type (color-to-shape, shape-to-color). Top row, monkey W; bottom row, monkey J. **c,** Proportion of trials that monkeys aborted; conventions as for panel b. **d,** Accuracy on each trial type as a function of the number of times that trial type was presented; there are 12 trial types for each set of object concepts (six object concepts, 2 trial types per concept: color-to-shape, shape-to-color).

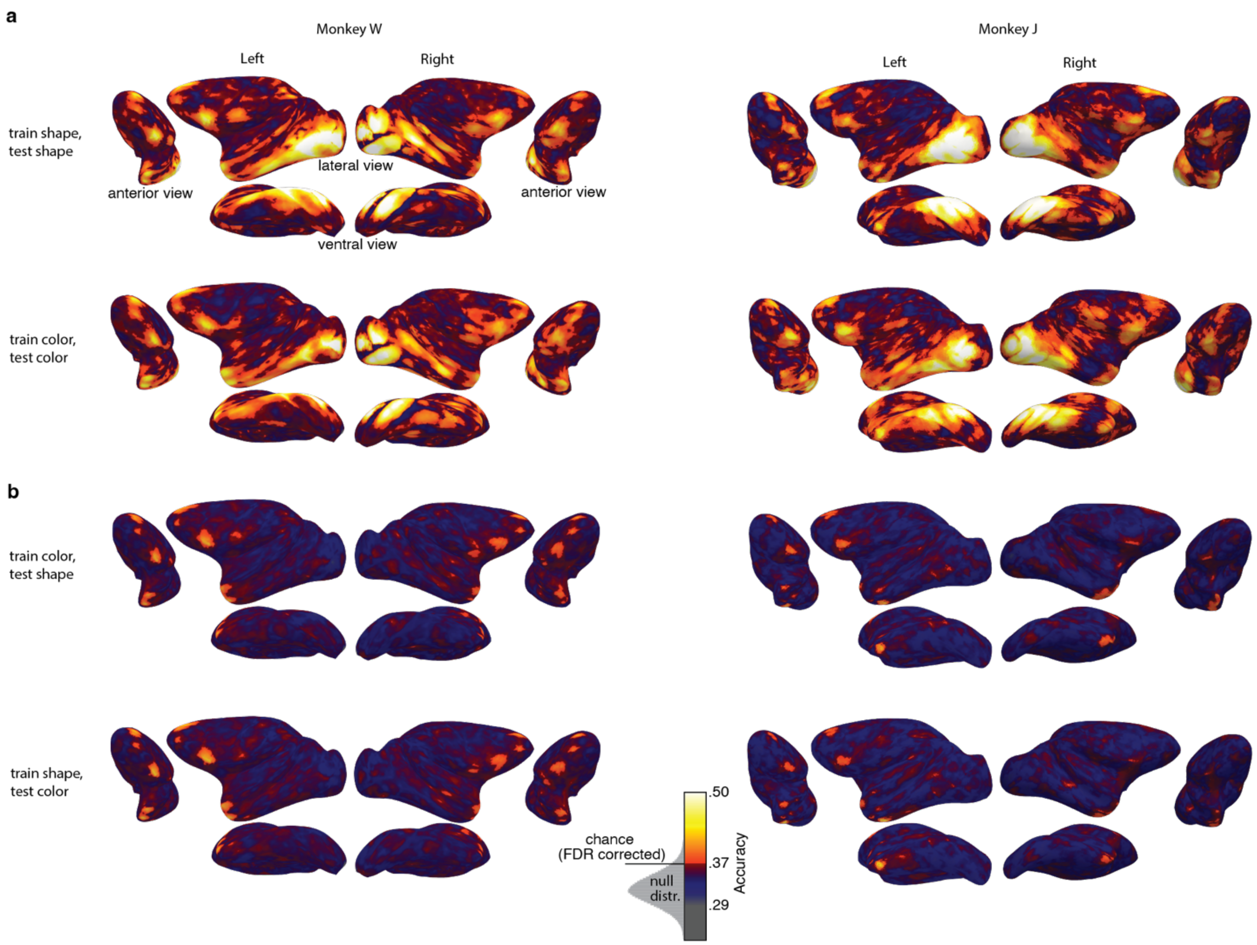


**Fig. S3. Cortical surface maps showing object-concept feature (color, shape) identity decoding and cross-feature decoding of fMRI data collected in the two macaque monkeys performing long-term memory trials (see Fig. 2a for trial structure). a,** Shape-identity decoding (train shape, test shape) and color-identity decoding (train color, test color) for both cortical hemispheres of both monkeys, three views per hemisphere (from anterior, ventral, and lateral positions). **b,** Color-to-shape (train color, test shape) and shape-to-color (train shape, test color) cross decoding conventions as for panel a. Color-scale bar shows decoding accuracy and the null distribution used to determine the FDR-corrected chance threshold.

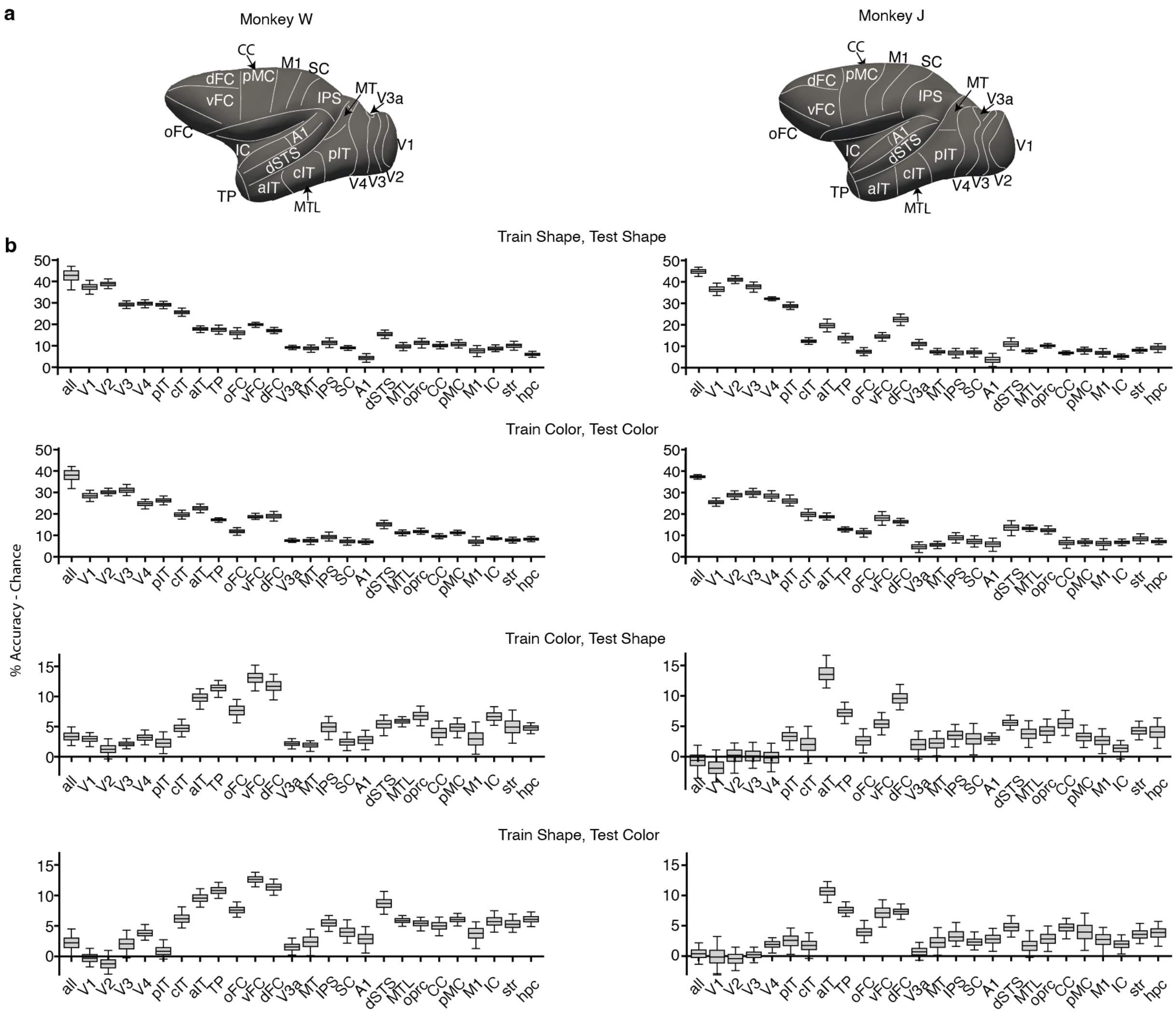


**Fig. S4 Object concept feature identity and cross-feature decoding quantified in brain parcels (see Fig. 2a for trial structure). a,** Brain parcels shown on the inflated cortical surface of the left hemisphere of each monkey. **b,** From top to bottom: shape-identity decoding, color-identity decoding, color-to-shape cross decoding, and shape-to-color cross decoding. Analysis was data-driven using the LSDM across the whole brain (see methods for description of the LSDM). The LSDM searchlight patches comprising a given brain parcel were aggregated to generate the box plots (see Methods). Brain parcels are: a grand parcel including all parcels (all); retinotopic cortical areas (V1 to V4); posterior inferior temporal cortex (pIT), central IT (cIT), anterior IT (aIT), temporal pole (TP), orbital frontal cortex (oFC), ventral frontal cortex (vFC), dorsal frontal cortex (dFC), dorsal visual areas (V3a, middle temporal area MT), intraparietal sulcus (IPS), somatosensory cortex (SC), primary auditory cortex (A1), dorsal superior temporal sulcus (dSTS), medial temporal lobe regions of entorhinal cortex and parahippocampal cortex (MTL), operculum region superior to the insula and inferior to motor cortex (oprc), cingulate cortex (CC), insula cortex (insl), striatum (str), and hippocampus (hpc). Boxes show percent accuracy above chance, data averaged across hemispheres and monkeys, error bars show 95% CI.

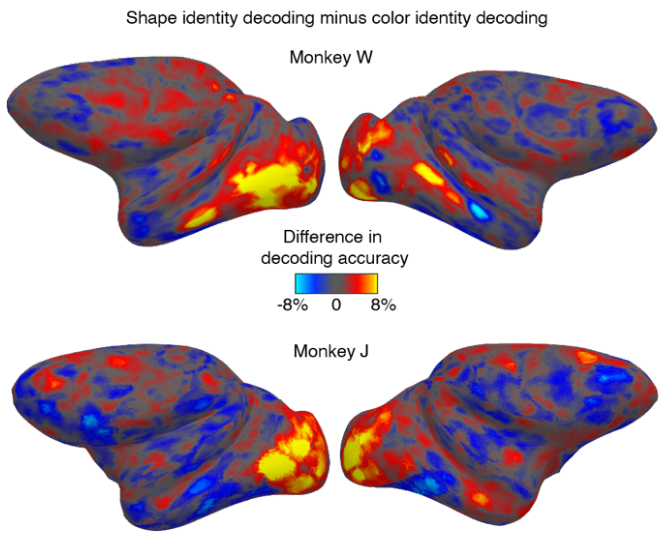


**Fig. S5 Comparison of shape-identity decoding and color-identity decoding of fMRI data collected while monkeys perform long-term memory trials (see Fig. 2a for trial structure).** The plots show surface maps of both cerebral hemispheres for the two monkeys, with bias for shape-identity decoding (warm colors) and color-identity decoding (cool colors). Color bar indicates the difference in accuracy between shape identity (train shape, test shape) decoding and color identity (train color, test color) decoding.

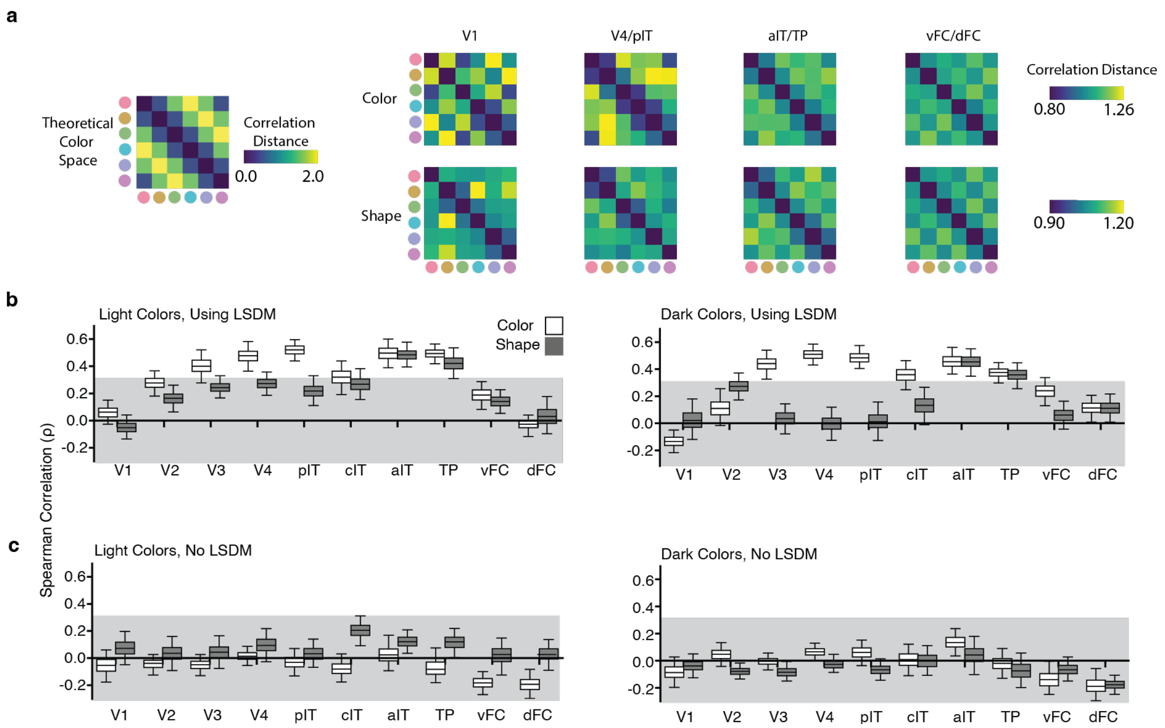


**Fig. S6 Structure of the neural representation of object concepts computed using fMRI data collected while monkeys performed long-term memory trials (see Fig. 2a for trial structure). a,** Representational Dissimilarity Matrices (showing 1 – Pearson correlation) for color (top row) and shape (bottom row) stimuli computed in the LSDM latent space, averaged over monkeys and luminance levels. The theoretical representational dissimilarity matrix for the stimuli in CIELUV color space is shown on the left; yellow indicates two colors are far from one another in perceptual color space. **b,** Correlation of the pattern of fMRI responses elicited by object colors with color-space geometry (white boxes) and by object shapes with color-space geometry (gray boxes), computed with the LSDM model, for light colors (left) and dark colors (right). **c,** Correlation computed without the LSDM model for light colors (left) and dark colors (right). Representations in (b) were computed in the input voxel space of each searchlight patch using a standard representational similarity analysis (RSA) searchlight. The shaded region indicates correlation levels that could be achieved by chance (95% interval)

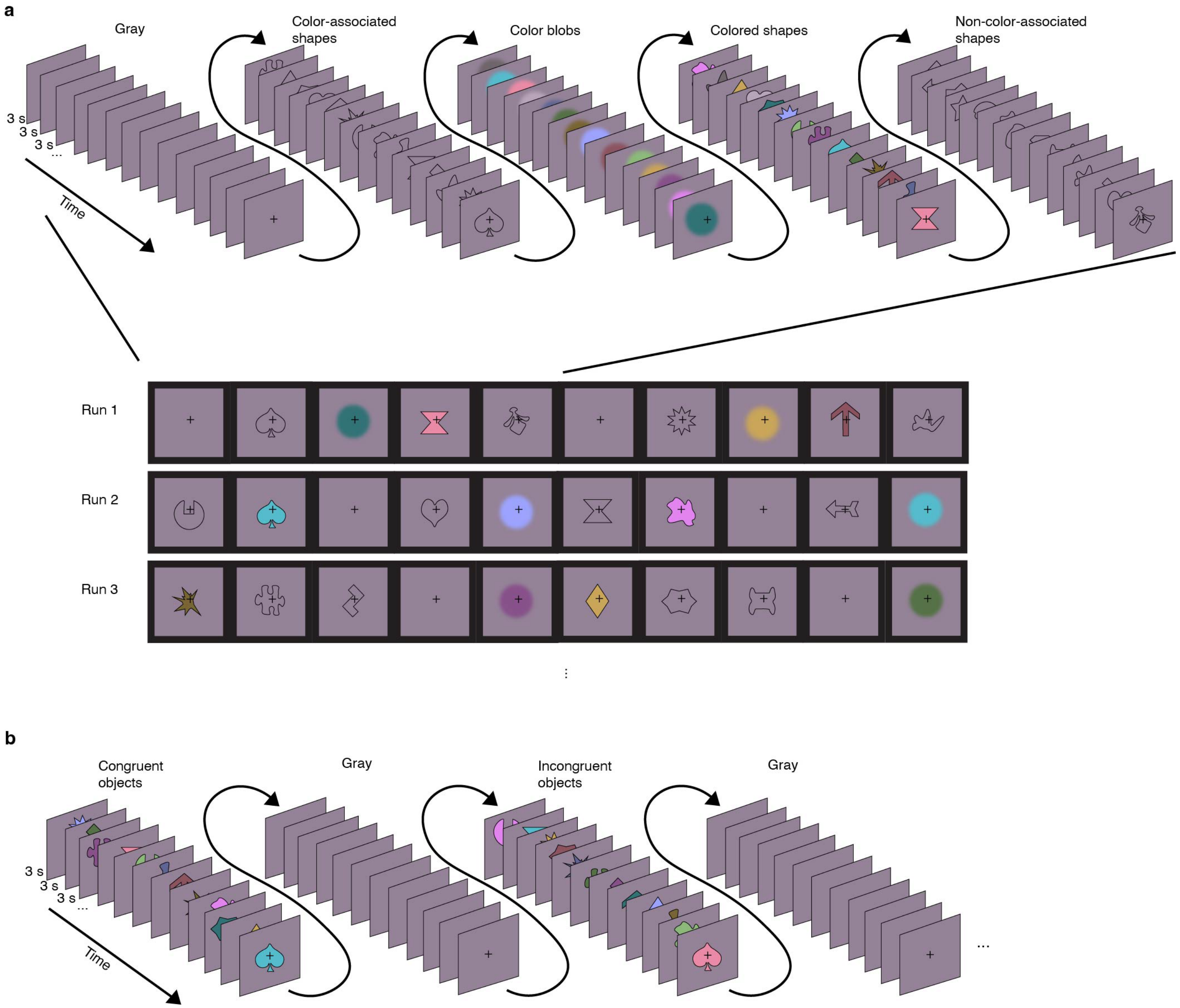


**Fig. S7 The paradigm for leveraging top-down feedback to identify areas responsible for (a) color perception and (b) color-shape binding. a,** Monkeys viewed blocks of colored shapes (the colored object concepts), shapeless color blobs (the colors of the colored object concepts), colorless color-associated shapes (the shapes of the colored object concepts), colorless non-color-associated shapes (colorless shape concepts), and full-field gray. In the present work, we only analyzed the responses to the two colorless shape conditions, which are matched in low-level stimulus features and differ because only for one of the conditions do the shapes have long-term associations with colors. Each block showed all stimuli from one condition once each in random order, and each stimulus was presented for 3 seconds (one TR) at the center of gaze with spatial jitter. A run consisted of ten blocks; the stimuli in the first five blocks of an example run are shown (top). The blocks were counterbalanced with deBruijn sequencing across runs. A fixation cross was presented at the center of the screen and the monkeys received juice reward at random times throughout the run for maintaining fixation. The first frame of each block for three example runs are shown (bottom). **b,** Monkeys viewed alternating blocks of colored shapes in which the color and shape of the stimuli were congruent (the color and shape of the stimulus agree with the learned association) or incongruent (the color and shape of the stimulus do not agree with the learned association). For example, the monkeys learned yellow-diamond and pink-hourglass concepts, these would be congruent stimuli, whereas a yellow-hourglass or pink-diamond would be incongruent stimuli. Each stimulus was presented for 3sec (one TR), and stimulus blocks were separated by gray blocks. Each block showed all shapes and all colors once each in random order. Each run contained 16 blocks. Other conventions as for the experiment in a.

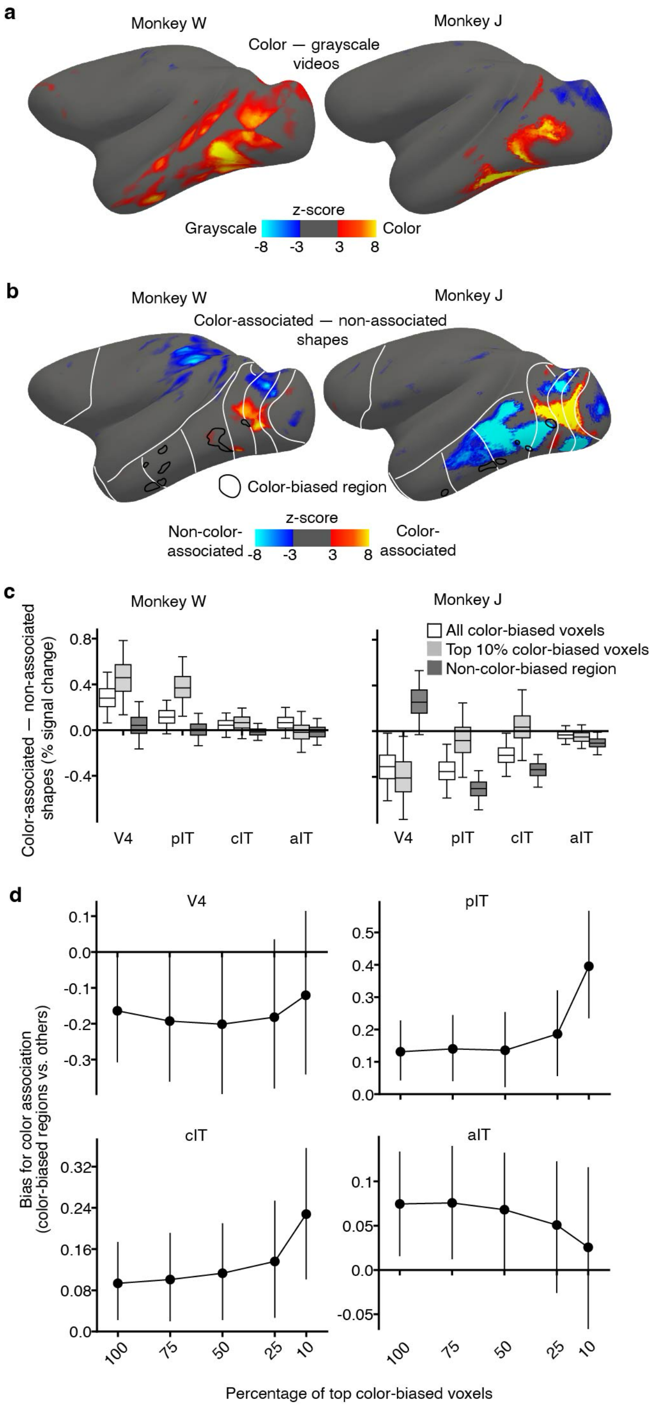


**Fig. S8. Do color-biased regions respond to object-color knowledge? a,** Differences in fMRI responses to videos of faces, bodies, scenes, objects, scrambled objects presented in color versus presented in grayscale across the cortical surface (Z-score contrast maps, bias for color in orange). **b,** Differences in fMRI responses to colorless color-associated shapes versus colorless non-color-associated shapes across the cortical surface, as shown in **Fig. 5a**, with color-biased regions outlined in black. Color-biased voxels were identified using the comparison shown in panel a. Color-biased regions outlined in black represent the 5% of voxels with the strongest color bias on the left hemisphere. **c,** Bias for colorless color-associated shapes versus colorless non-color-associated shapes (percent signal change difference), quantified in color-biased and non-color-biased regions across V4, pIT, cIT, and aIT. The proportion of voxels contributing to the analysis varied depending on their color bias (all color-biased voxels includes 100% of color-biased voxels at $p<.01$, one-sided, FDR-corrected; top 10% consists of 10% of voxels with the strongest color bias). The number of non-color-biased voxels was fixed, and determined as all visually driven voxels that were not color biased. Results are shown separately for the two monkeys. For Monkey W, most data points lie above zero, while for Monkey J most data points lie below zero. The relationship between color-biased regions and non-color-biased regions is consistent across monkeys in pIT, cIT, and aIT. **d,** Bias for colorless color-associated shapes compared to colorless non-color-associated shapes, for color-biased regions relative to non-color-biased regions in V4, pIT, cIT, and aIT, (direct comparison of the white and dark gray boxes in panel c, combined across monkeys, at different proportions of top color-biased voxels).

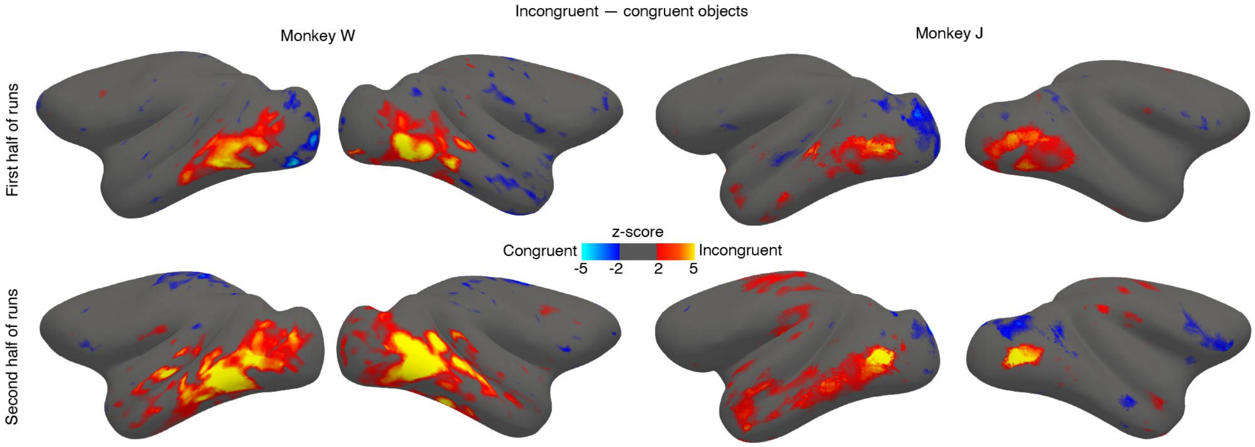


**Fig. S9. FMRI responses to incongruent versus congruent colored objects, for the first half and second half of fMRI runs collected in each monkey (see Fig. S7b for experimental paradigm).** Panels show cortical surface contrast maps, with the bias for incongruent colored objects shown in warm colors. Color scale bar shows z-scored difference between responses to incongruent objects and congruent objects using the beta weights assigned to each condition by the GLM. A total of 112 blocks per condition (across 4 sessions) for monkey W and 184 blocks per condition (across 5 sessions) for monkey J.

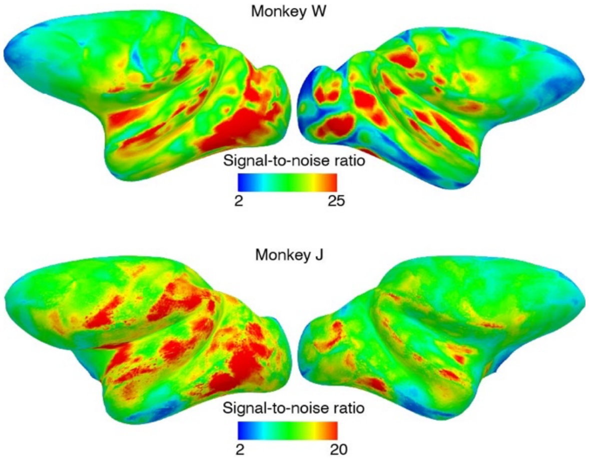


**Fig. S10 Signal-to-noise ratio (SNR) across the cortical surface of the two monkeys.** SNR was computed using all runs from the three main experiments (paradigms shown in **Fig. 2a, Fig. S7a, b**). For each voxel in the brain on each run, the voxel time course was averaged and divided by the average of the time course of a region consisting of 91 contiguous voxels outside of the brain. SNR was averaged across runs. Regions of lowest SNR, such as the temporal pole, were nonetheless of sufficient SNR to recover significant feature decoding (see **Fig. 2, S4**), implying that the variation in SNR across the cortical surface did not determine the pattern of decoding across the cortical surface.
